# Axin1 and Axin2 regulate the WNT-signaling landscape to promote distinct mesoderm programs

**DOI:** 10.1101/2024.09.11.612342

**Authors:** Rocío Hernández-Martínez, Sonja Nowotschin, Luke T.G. Harland, Ying-Yi Kuo, Bart Theeuwes, Berthold Göttgens, Elizabeth Lacy, Anna-Katerina Hadjantonakis, Kathryn V. Anderson

**Affiliations:** Developmental Biology Program, Sloan Kettering Institute, Memorial Sloan Kettering Cancer Center, New York, NY 10065, USA; Cambridge Stem Cell Institute, University of Cambridge, Cambridge, UK; Department of Haematology, University of Cambridge, Cambridge, UK

**Keywords:** Axin1, Axin2, WNT signaling, mouse embryo, gastrulation, mesoderm, extraembryonic, embryonic, BMP/pSMAD1/5/9, NODAL/pSMAD2/3

## Abstract

How distinct mesodermal lineages – extraembryonic, lateral, intermediate, paraxial and axial – are specified from pluripotent epiblast during gastrulation is a longstanding open question. By investigating AXIN, a negative regulator of the WNT/β-catenin pathway, we have uncovered new roles for WNT signaling in the determination of mesodermal fates. We undertook complementary approaches to dissect the role of WNT signaling that augmented a detailed analysis of *Axin1*;*Axin2* mutant mouse embryos, including single-cell and single-embryo transcriptomics, with *in vitro* pluripotent Epiblast-Like Cell differentiation assays. This strategy allowed us to reveal two layers of regulation. First, WNT initiates differentiation of primitive streak cells into mesoderm progenitors, and thereafter, WNT amplifies and cooperates with BMP/pSMAD1/5/9 or NODAL/pSMAD2/3 to propel differentiating mesoderm progenitors into either posterior streak derivatives or anterior streak derivatives, respectively. We propose that *Axin1* and *Axin2* prevent aberrant differentiation of pluripotent epiblast cells into mesoderm by spatially and temporally regulating WNT signaling levels.

## INTRODUCTION

Gastrulation converts pluripotent epiblast cells in a developing embryo into the three definitive germ layers: ectoderm, mesoderm, and endoderm. In the mouse, the emergence of the primitive streak in the proximal posterior epiblast marks the onset of gastrulation at embryonic day (E)6.25-6.5 (Bardot and Hadjantonakis, 2020). Epiblast cells residing in the vicinity of the primitive streak exit pluripotency and undergo an epithelial-to-mesenchymal transition (EMT) facilitating their exit from the streak and movement into the mesenchymal layer, where they migrate across the embryo and differentiate into progenitors of mesoderm and definitive endoderm (Ferrer-Vaquer et al., 2010a).

The signaling environment of epiblast cells at the time and site of their ingression through the primitive streak determines their fate, and thus the identity of the different mesodermal types produced (Morgani and Hadjantonakis, 2020). Fate mapping experiments in the mouse have shown that the first wave of mesoderm emanating from the streak includes progenitors of extraembryonic mesoderm (ExM) (Lawson et al., 1991); these cells migrate proximally to contribute to the amnion, allantois, chorion, and visceral yolk sac (Kinder et al., 1999; Lawson et al., 1991). Concurrently, from a slightly more distal location in the streak, the newly specified precursors of cardiac and cranial mesoderm migrate anteriorly. As the anterior primitive streak extends toward the distal tip of the embryo, it gives rise to paraxial and axial mesoderm (prechordal plate, notochord and node), as well as precursors of the definitive endoderm (Kinder et al., 1999; Kinder et al., 2001). Additional mesoderm subtypes, such as lateral plate, and intermediate mesoderm are generated from the middle primitive streak (Kinder et al., 1999; Lawson et al., 1991; Tam and Beddington, 1987).

Over the past three decades, genetic studies in the mouse have established that Bone Morphogenetic Proteins (BMPs) and NODAL, both TGF-ß family members and two of many diffusible ligand signaling molecules present in the embryo at these stages, play pivotal roles in regulating axis formation at the onset of gastrulation (Arnold and Robertson, 2009). In addition, analyses of mouse embryos with altered alleles of individual BMP and NODAL pathway components, suggest that regulatory feedback loops between NODAL and BMP operate in the streak to confer cell fate (Ben-Haim et al., 2006). For example, embryos lacking *Bmpr1a* (Di-Gregorio et al., 2007; Mishina et al., 1995) or *Nodal* (Brennan et al., 2001) fail to establish a streak and to make mesoderm. Embryos mutant for *Bmp4* (Winnier et al., 1995) or for the SMAD1/5/9 downstream components, display defective formation of the allantois and yolk sac, suggesting that specification of ExM requires a functional BMP/pSMAD1/5/9 signaling pathway (Chang et al., 1999; Tremblay et al., 2001). In contrast, mouse embryos with a combined loss of NODAL intracellular effectors, *Smad2* and *Smad3,* in the epiblast, generate an enlarged allantois, but lack the axial, paraxial and lateral populations of embryonic mesoderm (Dunn et al., 2004). However, it remains unclear how BMP and NODAL signals coordinate with one another and are combinatorially interpreted by cells *in vivo* to generate spatially and functionally distinct populations of mesodermal cell types.

Acting through its effector β-CATENIN, canonical WNT signaling also plays a critical role in multiple developmental events in the early mouse embryo, including axis formation and cell fate specification. Embryos mutant in *Wnt3* fail to gastrulate and thus lack mesoderm (Liu et al., 1999). In contrast, in mouse mutants unable to down-regulate β-CATENIN effectors, such as *APC^min^* (Chazaud and Rossant, 2006) and β*-catenin*^Δ*exon3*^ (Kemler et al., 2004), an excess of WNT signaling drives the epiblast to prematurely exit pluripotency and express mesenchymal markers. These findings suggest that modulation of WNT signaling levels in epiblast cells during their exit from pluripotency achieves activation of stage and region-specific targets that contribute to the acquisition of distinct cell fates. AXIN1 and AXIN2 are components of the destruction complex, which inhibits canonical WNT signaling through the phosphorylation, ubiquitination, and degradation of β-CATENIN (Ikeda et al., 1998). Loss of function alleles of *Axin1* cause embryonic lethality: mutant mouse embryos display multiple morphological abnormalities, including cardia bifida and partial duplications of the body axis, and they die at around E9.0 (Perry et al., 1995; Zeng et al., 1997a). In contrast, *Axin2* null mutants are viable and fertile (Lustig et al., 2001a). AXIN2 protein is structurally very similar to AXIN1, and lack of *Axin1* can be functionally replaced by *Axin2 in vivo* (Chia and Costantini, 2005).

The dose and time-dependent mechanisms by which WNT interacts with other signals at the primitive streak to confer mesodermal identity *in vivo* remain unclear. To ask if the WNT pathway interfaces with the NODAL and BMP pathways *in vivo* to direct the specification of mesodermal lineages, we examined mouse embryos carrying combinations of mutations in key components of the WNT, BMP and NODAL signaling pathways. First, focusing on *Axin1;Axin2* double mutants, we show that constitutively active WNT signaling drives the early post-implantation epiblast to prematurely transition from the formative to the primed state of pluripotency and initiate an EMT. Second, using *Axin1;Axin2* double mutants lacking *Axin1* exclusively in the epiblast (*Axin1*^Δ*Epi*^*;Axin2^-/-^*), we demonstrate that constitutively active WNT signaling propels the epiblast towards differentiation into posterior and middle streak mesodermal lineages at the expense of anterior streak mesodermal derivatives. To determine how constitutively active WNT signaling disrupts the coordinated production of mesoderm at the streak, we performed both single-cell RNAseq (scRNAseq) and single-embryo bulk RNAseq analyses in E6.5 (at the onset of gastrulation) and E7.5 *Axin1*^Δ*Epi*^*;Axin2^-/-^*mutants. Our analysis of these transcriptomic data reveal that high levels of BMP/pSMAD1/5/9 accompany aberrant posterior mesoderm differentiation in both the *Axin1*^Δ*Epi*^*;Axin2^-/-^* and *Smad2^-/-^*mutants, suggesting that posterior mesodermal identity depends on coordinated activity of WNT and BMP, but is independent of NODAL signaling. To further dissect how the network of signaling interactions regulate the production of different mesodermal types, we performed *in vitro* pluripotent Epiblast-Like Cell (EpiLC) differentiation experiments. Taken together, we propose a working model positing that WNT signals prime epiblast cells to differentiate into mesodermal progenitors; concurrent BMP/pSMAD1/5/9 signals then direct these progenitors to form extraembryonic, posterior lateral plate and intermediate mesoderm, while anterior mesoderm derivatives require a tighter WNT-BMP spatial-temporal regulation to maintain NODAL signaling and give rise to axial and paraxial mesoderm.

## RESULTS

### Constitutively active WNT signaling precociously drives epiblast cells through formative pluripotency into a primed state before the onset of gastrulation

*Axin1* is ubiquitously expressed in the early embryo (Qian et al., 2011) and its loss causes mid-gestation embryonic lethality. In contrast, induction of *Axin2* expression occurs only upon activation by WNT (Jho et al., 2002), and targeted *Axin2* null alleles produce viable and fertile mice (Lustig et al., 2001a). Uniform Manifold Approximation and Projection (UMAP) plots from a publicly available scRNAseq reference atlas of mouse gastrulation (Imaz-Rosshandler et al., 2024; Pijuan-Sala et al., 2019) show that both *Axin1* and *Axin2* are widely expressed. *Axin2* shows expression enrichment in epiblast, anterior primitive streak and mesoderm cells (**Figure 1A**; see **Figure S2A** for cell type reference). To assess the effects of different levels of elevated WNT signaling on epiblast maturation, and the initiation of gastrulation defined by pluripotency exit, we examined embryos carrying different combinations of *Axin1* and *Axin2* null alleles (**Figure 1B**). Although recoverable at early gastrulation (E6.25), embryos deficient for both *Axin1* and *Axin2* (*Axin1^-/-^;Axin2^-/-^; Axin1^fus/fus^; Axin2^-/-^)* were morphologically distinct from stage-matched wild-type embryos. In addition, immunofluorescence (IF) analyses on E5.75 *Axin1;Axin2* null mutants revealed that the entire epiblast had prematurely differentiated into mesoderm with up-regulated T (a marker of the primitive streak and nascent mesoderm) and down-regulated CDH1 expression (a marker downregulated during the gastrulation EMT, **Figure 1C** and **Figure S1A**). Embryos lacking *Axin1* specifically in epiblast cells and their derivatives but containing two wild-type *Axin2* alleles in all cells (*Axin1^ΔEpi^*;*Axin2^+/+^*), developed to E9.5 displaying defects in the formation of forebrain, heart, somites, and tail among other developing tissues (**Figure S1B**). At E5.75, both *Axin1^ΔEpi^*;*Axin2^+/+^*and *Axin1^ΔEpi^;Axin2^-/-^*mutant embryos appeared morphologically indistinguishable from wild-type embryos. However, IF of T and CDH1 revealed that loss of both alleles of *Axin2* in *Axin1^ΔEpi^*embryos (*Axin1^ΔEpi^;Axin2^-/-^*) resulted in a more severe phenotype than observed in *Axin1^ΔEpi^*;*Axin2^+/+^* embryos. While *Axin1^ΔEpi^*;*Axin2^+/+^*embryos did not exhibit detectable T expression or premature activation of EMT, *Axin1^ΔEpi^;Axin2^-/-^*mutants displayed ectopic T-expressing mesoderm cells at E5.75, before E6.25-E6.5, the age at which initiation of gastrulation normally occurs (**Figure 1C**). A comparison of the CDH1 and T staining patterns at E5.75 in *Axin1^-/-^*;*Axin2^-/-^* and *Axin1^ΔEpi^;Axin2^-/-^* embryos suggested a correlative relationship between the level and/or distribution of constitutively active WNT signaling and the ectopic mesoderm. For example, *Axin1^ΔEpi^;Axin2^-/-^* embryos retained the WNT inhibitory activities of AXIN1 that were lost in the visceral endoderm and extraembryonic ectoderm of *Axin1^-/-^;Axin2^-/-^* embryos. At E5.75, the epiblast in *Axin1^ΔEpi^;Axin2^-/-^* embryos contained fewer T-expressing cells and maintained a more organized epithelial structure than the epiblast in *Axin1^-/-^;Axin2^-/-^* embryos (**Figure 1C,D**).

**Figure 1.**
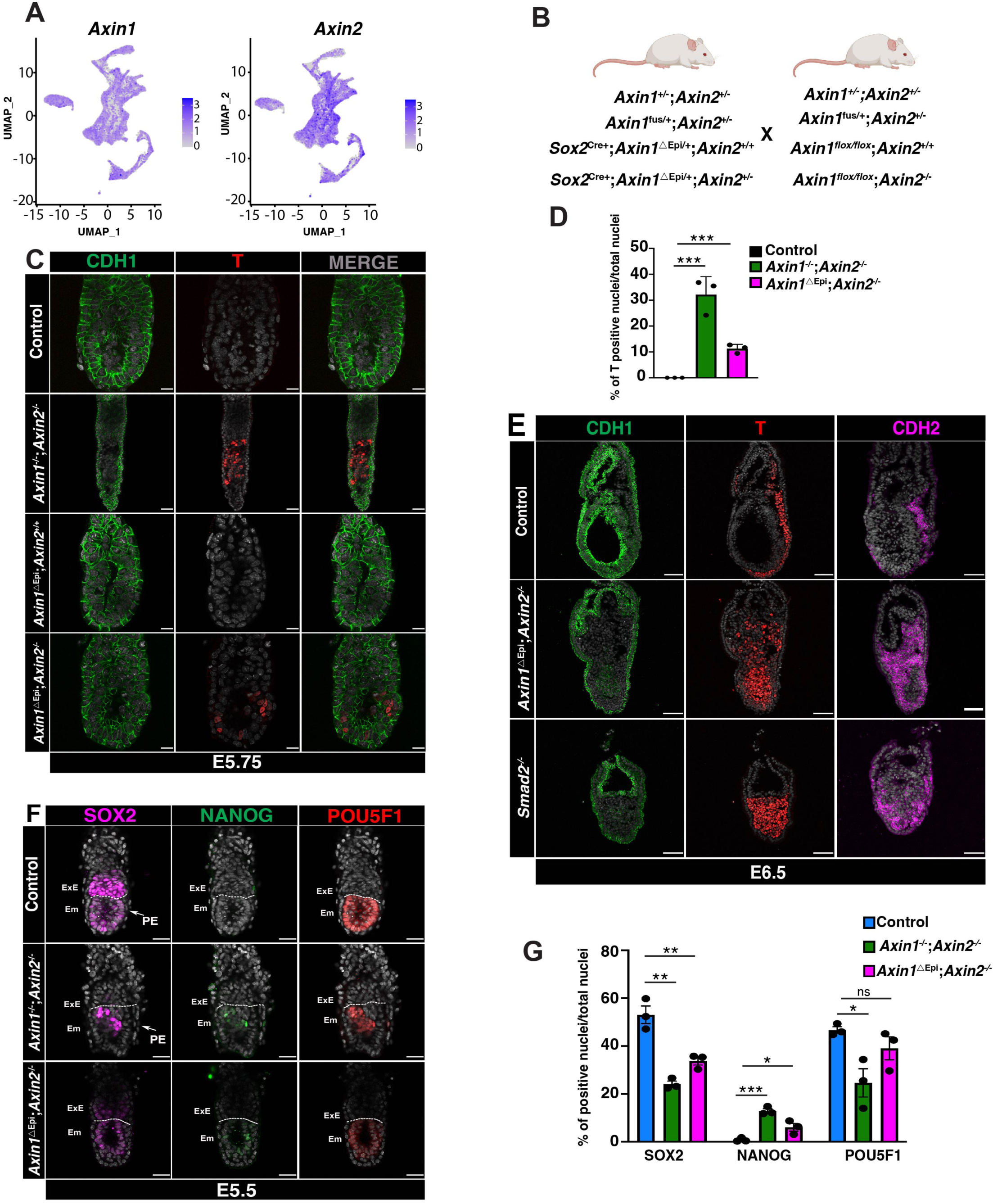
*Axin1* and *Axin2* prevent ectopic EMT and preserve epiblast naïve pluripotency. (**A**) UMAP obtained from the extended gastrulation mouse atlas showing *Axin1* and *Axin2* expression in the single cell lineages. (**B**) Cartoon showing the breeding strategy. “Created with BioRender.com”. (**C**) Single optical sections of whole-mount immunofluorescence (IF) detecting T and CDH1 in E5.75 wild-type, *Axin1^-/-^;Axin2^-/-^*and *Axin1^ΔEpi^;Axin2* mutant embryos. (**D**) Graph showing percentage ratio of T-positive nuclei relative to the total nuclei in E5.75 wild-type, *Axin1^-/-^;Axin2^-/-^*and *Axin1^ΔEpi^;Axin2^-/-^* embryos. (**E**) IF on E6.5 longitudinal sections for CDH1, T and CDH2 in wild-type, *Axin1^ΔEpi^;Axin2^-/-^* and *Smad2^-/-^*embryos. (**F**) Single optical sections of whole-mount IF detecting SOX2, NANOG and POU5F1 in E5.5 wild-type, *Axin1^-/-^;Axin2^-/-^*and *Axin1^ΔEpi^;Axin2^-/-^*embryos. PE (Primitive Endoderm), ExE (Extraembryonic Ectoderm) and Em (embryo). Scale bars are 50μm. (**G**) Graph showing the ratio of positive nuclei for the different markers relative to the total nuclei in the E5.5 wild-type, *Axin1^-/-^;Axin2^-/-^*and *Axin1^ΔEpi^;Axin2^-/-^*embryos. Bars in **D** and **G** represent the mean± SEM. Statistically significant differences were determined by t-test. *p<0.03, **p<0.009, ***p<0.0003, ns=not significant.

Examination of *Axin1^ΔEpi^;Axin2^-/-^* embryos at E6.5 provided further evidence that epiblast cells had undergone a premature, delocalized EMT program. During gastrulation EMT, nascent mesoderm cells emanating from the primitive streak downregulate CDH1, upregulate CDH2, and activate expression of T (Hernández-Martínez et al., 2019), generating a posteriorly localized stripe of CDH2 and T-expressing cells by E6.5 (**Figure 1E**). In contrast, we detected CDH2 and T-expressing cells widely dispersed throughout the proximal and distal regions of the E6.5 *Axin1^ΔEpi^; Axin2^-/-^*embryos (**Figure 1E**).

To confirm that the formation of ectopic mesoderm reflected premature differentiation of the epiblast, we performed whole-mount IF of markers diagnostic for the naïve and primed pluripotent states of epiblast cells in pre-implantation and gastrulation stage embryos, respectively (Morgani et al., 2017; Nichols and Smith, 2009). Following implantation, but before gastrulation, epiblast cells reside in a transitional state, referred to as formative pluripotency (Smith, 2017), during which they down-regulate the naïve transcriptional program (POU5F1, NANOG, SOX2) before actively expressing lineage-associated genes (T, FOXA2, CDX2) (Morgani et al., 2017; Smith, 2017). Relative to levels observed in wild-type embryos, we detected reduced SOX2 and OCT4, and increased NANOG expression in the epiblast of E5.5 *Axin1^-/-^;Axin2^-/-^*and *Axin1^ΔEpi^;Axin2^-/-^*mutants (**Figure 1F-G**). Using IF we found reduced expression of all three markers in E6.5 *Axin1^ΔEpi^;Axin2^-/-^* embryos (**Figure S1C**). Together these data show that in *Axin1/2* double mutant embryos, the epiblast undergoes premature down-regulation of pluripotency markers concomitantly with the formation of ectopic T-expressing mesoderm. Thus, constitutively active WNT signaling, resulting from loss of the AXIN1 and AXIN2 inhibitors, precociously drives epiblast cells through formative pluripotency into a primed state at E5.5, almost 24hr before the onset of gastrulation in wild-type embryos.

Similar to our findings in *Axin1^ΔEpi^;Axin2^-/-^*mutants, published studies have reported an expanded region of primitive streak marker (*T, Fgf8*) expression in early post-implantation embryos lacking *Smad2* (Waldrip et al., 1998). To directly compare the gastrulation defects observed in *Smad2* null embryos (see Methods) to the abnormalities in mesoderm formation in *Axin1^ΔEpi^;Axin2^-/-^* embryos, we performed IF staining of *Smad2^-/-^* mutants for markers of EMT and the primitive streak. In agreement with previous reports, we detected ectopic T expression throughout the epiblast, in about 53% of the cells comprising the E6.5 *Smad2^-/-^* embryo (**Figure 1E and Figure S1D**). Suggestive of a premature EMT analogous to that displayed in *Axin1^ΔEpi^;Axin2^-/-^* embryos, we observed the downregulation of CDH1 and upregulation of CDH2 in the epiblast of *Smad2* mutants (**Figure 1E**). We also found reduced expression of pluripotency-associated markers SOX2, NANOG and OCT4, in E6.5 *Smad2^-/-^* embryos (**Figure S1C**). However, in contrast to the E5.75 *Axin1^ΔEpi^;Axin2^-/-^* embryo, the E5.75 *Smad2^-/-^* mutant lacked detectable ectopic T-expression (**Figure S1A**). This finding suggests that at pre-gastrulation stages, abnormally elevated levels of WNT, but not reduced levels of NODAL signaling, can trigger the epiblast to undergo premature EMT and mesoderm production.

### Constitutively active WNT promotes ectopic extraembryonic mesoderm (ExM)/posterior lateral plate mesoderm (LPM) differentiation in the gastrulating mouse embryo

To explore shifts in cellular heterogeneity and changes in transcriptional profiles, driven by constitutively active WNT signaling, we collected separate pools of control and *Axin1^ΔEpi^;Axin2^-/-^*double mutant mouse embryos at E6.5 and performed scRNAseq (**Figure 2**). In total, 10,250 single-cell transcriptional profiles passed quality control; these profiles were placed into a developmental context by integrating our datasets with a time-resolved scRNAseq reference atlas of mouse gastrulation that spans E6.5 to E8.5 (Imaz-Rosshandler et al., 2024; Pijuan-Sala et al., 2019)(**Figure S2A-B**). After integration, we investigated marker gene expression patterns, performed joint clustering, and transferred cell type and embryonic stage labels from the reference atlas to our query datasets (**Figure 2A-G**). Altogether, utilizing this information, we assigned cell type annotations to 16 joint clusters that were present in the control and *Axin1^ΔEpi^;Axin2^-/-^*embryos, as well as the reference atlas (**Figure 2A-G** and **Figure S2A**-**C**).

**Figure 2.**
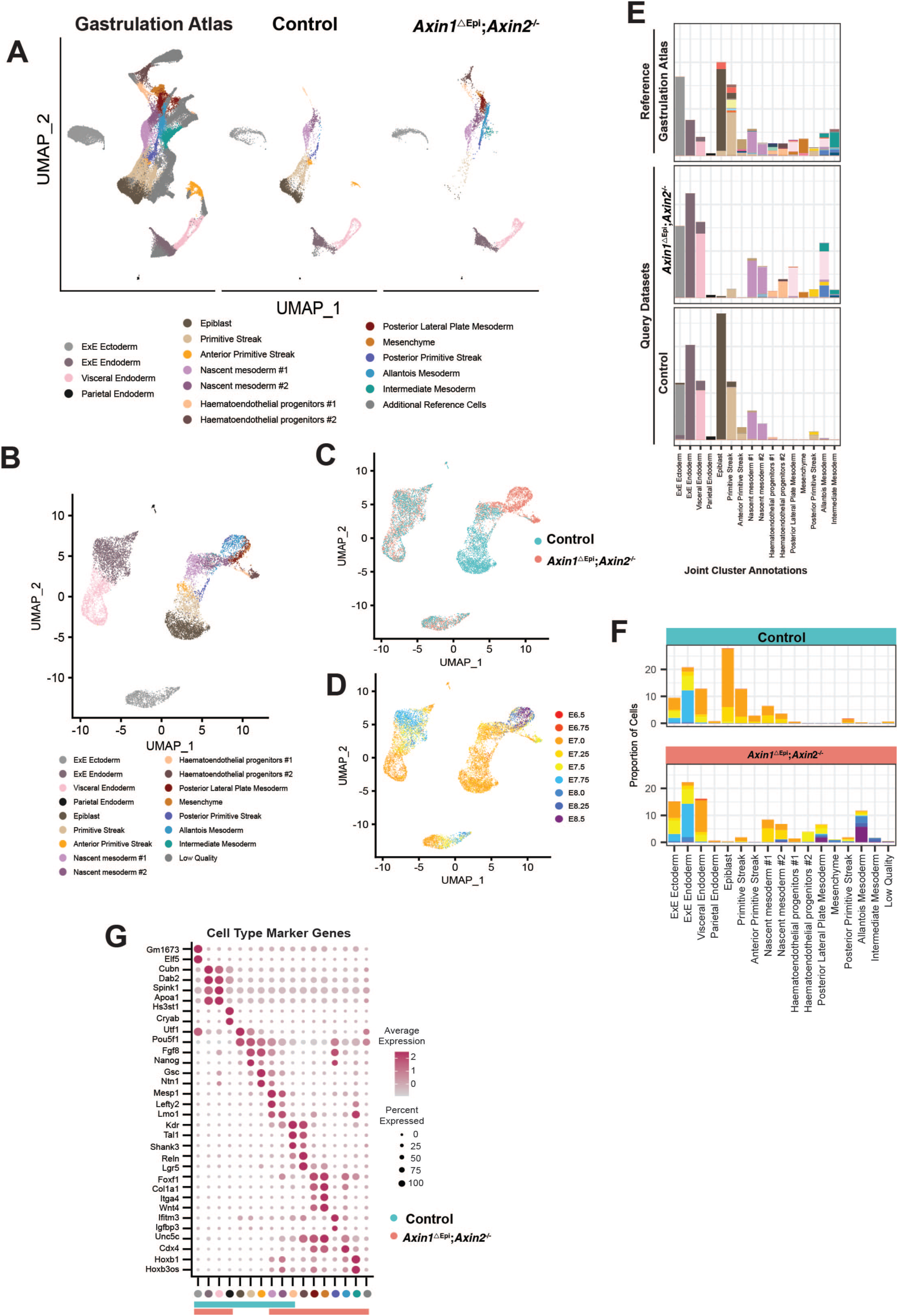
*Axin1* and *Axin2* prevent aberrant extraembryonic mesoderm, posterior lateral plate, and intermediate mesoderm differentiation. **(A)** UMAP generated using integrated scRNA-seq expression data from E6.5 control and *Axin1^ΔEpi^;Axin2^-/-^* embryos and a down sampled version of the gastrulation atlas that spans E6.5 to E8.5. Cells are colored by cell type annotations**. (B-D)** UMAP generated using integrated scRNA-seq expression data from E6.5 control and *Axin1^ΔEpi^;Axin2^-/-^* embryos. Cells are colored by cell type annotations in B, genotype in **C** and by embryonic stage. (**D**) Embryonic stage was transferred from the nearest neighbors from the mouse gastrulation atlas**. (E)** Stacked bar plots showing the proportion of different cell types in the gastrulation atlas (top), *Axin1^ΔEpi^;Axin2^-/-^* (middle) and Control (bottom) embryos. Bars are filled with the extended gastrulation atlas cell colors that indicate the proportion of cells within each cell type that are assigned different extended atlas cell type stage labels. In the Control (bottom) and *Axin1^ΔEpi^;Axin2^-/-^*(middle) bar plots, the extended atlas cell type labels were assigned via label transfer from the nearest neighbors in the gastrulation reference atlas. **(F)** Stacked bar plot showing the proportion of different cell types in Control (top) and *Axin1^ΔEpi^;Axin2^-/-^* (bottom) embryos. Bars are filled with embryonic stage colors that indicate the proportion of cells within each cell type that assigned different embryonic stage labels via label transfer from the nearest neighbors in the gastrulation reference atlas. **(G)** Dot plot showing log normalized expression levels of top marker genes (avg_log2FC > 0.5) for the cell types show in **B**. The size of the dot indicates the percentage of cells expressing the marker gene.

Approximately 50% of cells in control embryos aligned transcriptionally with epiblast (*Pou5f1*) or epiblast-derived cell types that predominate in E7.0-E7.25 embryos, including primitive streak (*Nanog, Fgf8*), anterior primitive streak (*Gsc*), posterior primitive streak (*Ifitm3, Igfbp3*), nascent mesoderm #1/#2 (*Mesp1, Lefty2*) and hematoendothelial progenitors #1 (*Kdr,Tal1*) (**Figure 2E-G**). The remaining control cells (∼50%) were given trophectoderm and primitive endoderm derived lineage annotations, including extraembryonic ectoderm (ExE), (*Elf5)*, extraembryonic endoderm (ExEn)/visceral endoderm (*Ttr, Spink1*), and parietal endoderm (PE) (*Cryab, Nid1*) (**Figure 2E-G**).

Like control embryos, approximately 50% of cells in *Axin1^ΔEpi^;Axin2^-/-^*mutant embryos aligned with ExE, ExEn/visceral endoderm, and PE (**Figure 2E-G**). Strikingly, instead of comprising the epiblast, primitive streak and nascent mesodermal cells, the remaining cells in *Axin1^ΔEpi^;Axin2^-/-^*mutant embryos represented more mature mesoderm lineages that normally only arise at later embryonic stages (E7.5-E8.5) (**Figure 2E-G**). Approximately 20% of cells in mutant embryos were posterior LPM, allantois mesoderm and mesenchymal cells that expressed *Foxf1*, *Cdx2*, *Cdx4* and *Twist2* (**Figure 2E-G and Figure S2C**). Furthermore, mutant embryos contained hematoendothelial progenitors #1 and further differentiated hematoendothelial progenitors #2 (∼5%) that expressed primitive erythrocyte markers *Gata1, Runx1, Hbb-bh1* and *Hbb-y* (**Figure 2E-G**). Overall, the integration of control and *Axin1^ΔEpi^;Axin2^-/-^* data sets with the time-resolved scRNAseq reference atlas of mouse gastrulation reveals that constitutively active WNT signaling rapidly drives epiblast cells to precociously differentiate into mesodermal components of extraembryonic tissues, such as visceral yolk sac and allantois. A comparative analysis of single-embryo bulk RNAseq data from four control and four *Axin1^ΔEpi^;Axin2^-/-^* mutant embryos at E6.5 yielded findings in agreement with those from our scRNAseq studies (**Figure S2D**-**F**). Evaluation of the bulk RNAseq data identified 2373 upregulated genes and 1741 downregulated genes (fold change p<0.05) (**Table S1-S2**). Predominant among the upregulated genes were those with known developmental roles associated with the differentiation of ExM, such as primitive blood precursors (i.e.*Tal1*, *Cd44*, *Runx1*) (Ferkowicz et al., 2003; Shivdasanl et al., 1995; Yokomizo et al., 2008) (**Figure S2E** and **Table S1)** and of the allantois (i.e. *Tbx4*, *Kdr*, *Fgf18*, *Cdx2*) (Boylan et al., 2020; Drake and Fleming, 2000; Naiche and Papaioannou, 2003) (**Figure S2F** and **Table S1**).

### WNT signals modulate BMP/pSMAD1/5/9 in the proximal epiblast to trigger ExM and posterior LMP differentiation

The proportion of cells expressing ExM/posterior LPM markers *Kdr*, *Cdx2* and *Foxf1* increased in *Axin1^ΔEpi^;Axin2^-/-^*versus control embryos (**Figure 3A**). To validate this finding from our scRNAseq data, we assessed KDR, CDX2 and FOXF1 protein expression patterns in E6.5 wild-type and *Axin1^ΔEpi^;Axin2^-/-^* embryos (**Figure 3B**). In wild-type embryos about 6% of total cells expressed KDR in the proximal posterior portion of the embryonic region. Furthermore, CDX2 expressing cells (12% of total) resided predominantly in the ExE, whereas FOXF1 expressing cells (< 1% of the total) inhabited the proximal posterior portion of the embryonic region (**Figure 3B** and **Figure S3A**). In contrast, in E6.5 *Axin1^ΔEpi^;Axin2^-/-^* embryos, we detected an expanded region of KDR expression, comprising about 22% of total cells alongside ectopically expressed CDX2 (55%) and FOXF1 (39%) (**Figure 3B** and **Figure S3A**). IF to KDR, CDX2 and FOXF1 showed that the ectopic mesoderm present in E6.5 *Smad2* mutants also had an ExM identity (**Figure 3B**). Compared to *Axin1^ΔEpi^;Axin2^-/-^*embryos, the *Smad2* mutants differed in the spatial distribution and relative numbers of KDR, CDX2 and FOXF1 (38%, 59% and 13%) respectively (**Figure S3A**). Yet the findings of a preponderance of ExM in E6.5 *Smad2^-/-^*mutants indicated that decreased levels of SMAD2-mediated NODAL signaling resulted in phenotypic features similar to those observed in *Axin1^ΔEpi^*;*Axin2^-/-^*embryos with constitutively active WNT signaling: premature ectopic differentiation of the epiblast layer into ExM.

**Figure 3.**
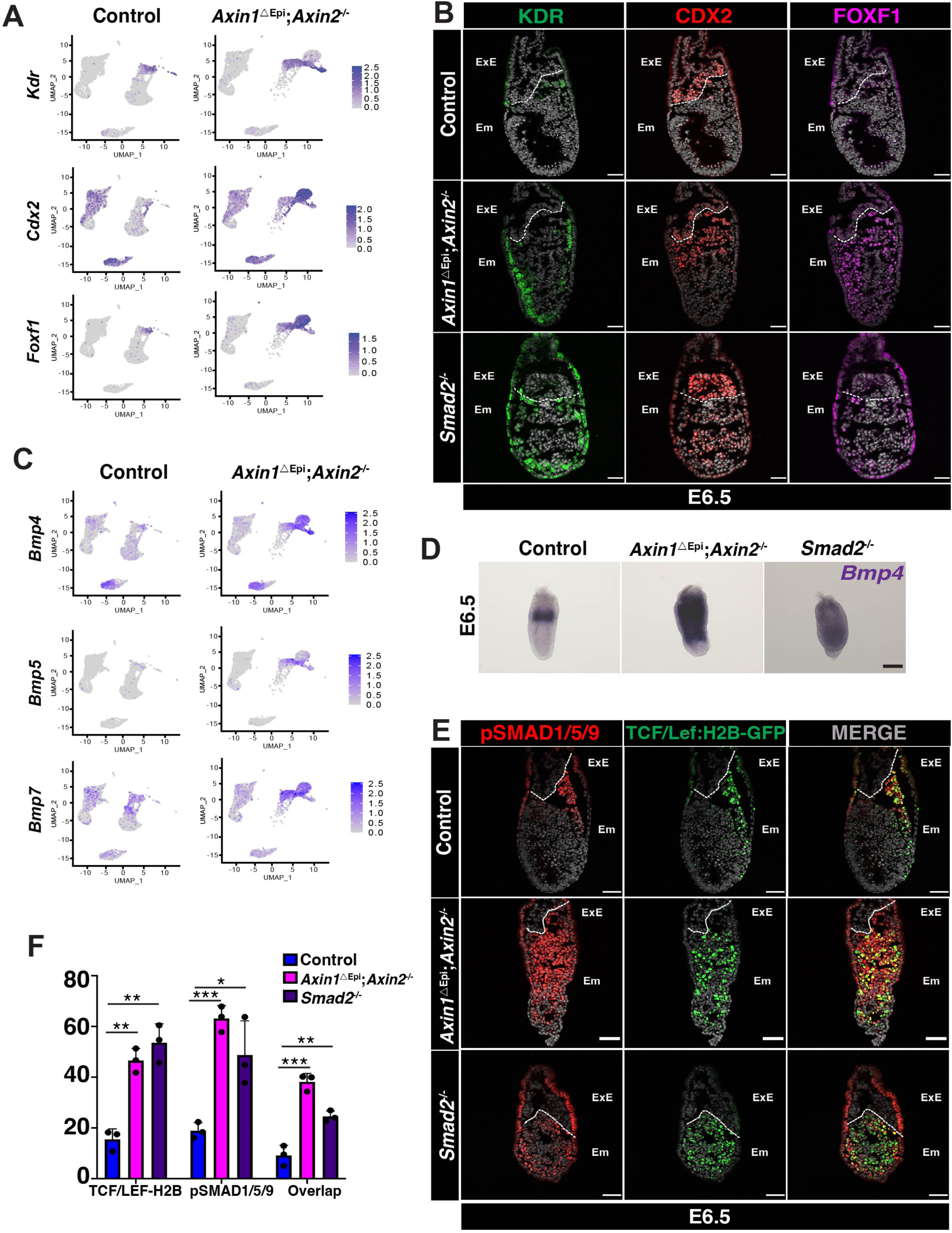
WNT signals trigger ExM differentiation by modulating BMP/pSMAD1/5/9 in the proximal epiblast. (**A**) UMAP generated using integrated scRNA-seq expression data, split by genotype, and colored by normalized gene expression of *Kdr* (top), *Cdx2* (middle) and *Foxf1* (bottom) in E6.5 wildtype and *Axin1^ΔEpi^;Axin2^-/-^* embryos. (**B**) IF staining of longitudinal cryosections of E6.5 wild-type, *Axin1^ΔEpi^;Axin2^-/-^* and *Smad2^-/-^*embryos detecting KDR, CDX2 and FOXF1. (**C**) UMAP generated using integrated scRNA-seq expression data, split by genotype, and colored by normalized gene expression of *Bmp4* (top), *Bmp5* (middle) and *Bmp7* (bottom) in E6.5 wildtype and *Axin1^ΔEpi^;Axin2^-/-^* embryos. (**D**) Whole-mount *in situ* hybridization detecting *Bmp4* expression in E6.5 control, *Axin1^ΔEpi^;Axin2^-/-^*and *Smad2^-/-^* embryos. Scale bar is 100μm. (**E**) IF in longitudinal sections detecting pSMAD1/5/9 and GFP on E6.5 wild-type, *Axin1^ΔEpi^;Axin2* and *Smad2* mutant. Scale bars on IFs are 50μm. (**B,E)** ExE (Extraembryonic Ectoderm) and Em (Embryo). (**F**) Graph showing ratio of GFP (TCF/Lef:H2B); pSMAD1/5/9 and nuclei positive for both GFP and pSMAD1/5/9 relative to the total of nuclei in the E6.5 wild-type, *Axin1^ΔEpi^;Axin2^-/-^* and *Smad2^-/-^* embryos. Bars represent the mean± SEM. Statistically significant differences were determined by t-test *p<0.02, **p<0.004, ***p<0.0002.

We examined the scRNAseq and single-embryo (se) bulk RNAseq data sets to ask if constitutively active WNT signals perturbed the expression of ligands in other key regulatory pathways active during early post-implantation development. This analysis revealed a strong upregulation of genes encoding the BMP ligands - *Bmp5*, *Bmp7, Bmp4* and *Bmp2* - in E6.5 *Axin1^ΔEpi^;Axin2^-/-^*mutants (**Figure 3C, Figure S3B** and **Table S1**). Using mRNA *in situ* hybridization, we confirmed the increased expression of *Bmp4* in E6.5 *Axin1^ΔEpi^;Axin2^-/-^*embryos. Whereas wild-type embryos contained *Bmp4* transcripts exclusively in the ExE, the *Axin1^ΔEpi^;Axin2^-/-^* and *Smad2^-/-^*mutants displayed an expanded domain of *Bmp4* expression that encompassed all but the distal portion of the epiblast (**Figure 3D**).

Elevated levels of *Bmp* gene expression upon constitutive WNT activity in *Axin1^ΔEpi^;Axin2^-/-^* embryos suggested that increased levels of BMP signaling may underlie the excess production of ExM/posterior LPM. To investigate a functional interaction between WNT and BMP signaling during gastrulation, we examined the spatial distribution and levels of WNT and BMP pathway activity by IF. We used a *TCF/Lef:H2B-GFP* reporter (Ferrer-Vaquer et al., 2010b) to identify cells transducing WNT and an antibody to pSMAD1/5/9 to detect BMP activated cells in E6.5 wild-type and *Axin1^ΔEpi^;Axin2^-/-^*embryos during early gastrulation (**Figure 3E-F**). In E6.5 wild-type embryos, pSMAD1/5/9 staining was restricted to cells in the proximal extraembryonic visceral endoderm (ExVE) and proximal epiblast ∼18% of the total cells in the embryo. WNT reporter activity was detected along the length of the streak in about 15% of the total cells in the embryo. By contrast, about 63% and 47% of total cells in the E6.5 *Axin1^ΔEpi^;Axin2^-/-^*embryo stained for pSMAD1/5/9 and the WNT signaling reporter, respectively (**Figure 3E-F**). We also observed that the fraction of cells that co-expressed both markers increased to 38% in mutant embryos relative to wild-type, where only 9% of cells co-expressed both markers (**Figure 3E-F**). In addition, we observed that the domain of pSMAD1/5/9 in the ExVE was extended distally in the *Axin1^ΔEpi^;Axin2^-/-^*embryos (**Figure 3E-F**).

Noting that expanded WNT and BMP signaling accompanied ectopic ExM/posterior LPM differentiation in *Axin1^ΔEpi^;Axin2^-/-^*mutant embryos, we asked whether *Smad2* mutants, which also abnormally differentiate ExM, similarly displayed increased numbers of WNT and BMP activated cells. We found that about 54% and 49% of cells in the E6.5 *Smad2^-/-^*embryo ectopically activated, *TCF/Lef:H2B-GFP* and pSMAD1/5/9, respectively. Concordantly, co-expression of pSMAD1/5/9 and the WNT-reporter was upregulated in about 25% cells in *Smad2^-/-^*embryos (**Figure 3E-F**). Taken together these observations suggest that the BMP-WNT signaling circuit that promotes ectopic ExM/posterior LPM differentiation in *Axin1^ΔEpi^;Axin2^-/-^* embryos also operates in the *Smad2* mutant (**Figure 3E-F**).

### Excess of WNT signaling depletes anterior primitive streak cell progenitors

To further examine the affected lineages in *Axin1^ΔEpi^;Axin2^-/-^* mutant embryos we performed se-bulk RNAseq on five wild-type and five *Axin1^ΔEpi^;Axin2^-/-^* E7.5 (mid-streak) embryos (**Figure S4A**). We found 2472 genes downregulated and 2493 genes upregulated in the mutants relative to wild-type (p<0.05) (**Table S3** and **Table S4**). Among the downregulated GO-terms were somitogenesis, rostro-caudal specification and NODAL signaling. Consistent with the findings in Figure 2 and 3, upregulated GO-terms included endothelial and blood vessel differentiation, EMT and BMP signaling (**Figure S4B**). We performed expression assays for two highly downregulated genes in the se-bulk transcriptome of E7.5 *Axin1^ΔEpi^;Axin2^-/-^*embryos: *Tbx6* and *Mesp1*, markers of paraxial mesoderm and cardiac progenitors, respectively (Chapman et al., 1996; Saga et al., 1999) (**Table S4**). As shown in **Figure 4A**, neither *Axin1^ΔEpi^;Axin2^-/-^* or *Smad2^-/-^*E7.5 embryos expressed detectable levels of TBX6 and *Mesp1*. We also did not detect expression of the axial mesoderm marker FOXA2 at E6.5 in either mutant (**Figure 4A**). By E7.5 both *Axin1^ΔEpi^;Axin2^-/-^*and *Smad2^-/-^* embryos are mainly composed of KDR, CDX2 and FOXF1-positive cells detected by IF (**Figure S4C**).

**Figure 4.**
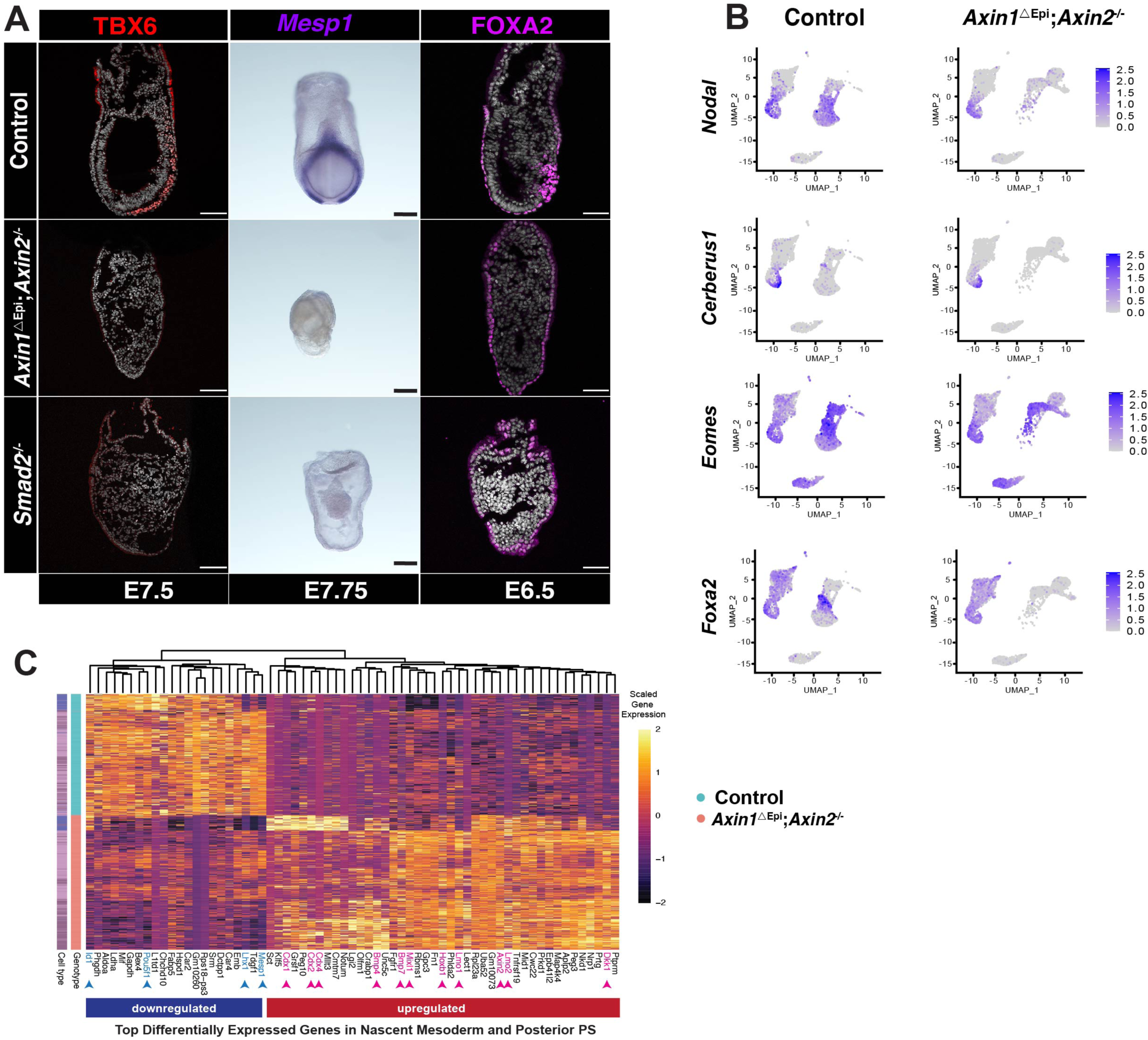
Excess of WNT activation blocks anterior primitive streak formation. (**A**) IF detecting TBX6 (E7.5) and FOXA2 (E6.5) on longitudinal sections of wild-type, *Axin1^ΔEpi^;Axin2^-/-^* and *Smad2^-/-^*embryos; whole-mount *in situ* hybridization to *Mesp1* (E7.75). Scale bar is 100μm. Scale bars on IFs are 50μm. (**B**) UMAP plots generated using integrated scRNA-seq expression data, split by genotype, and colored by normalized gene expression of *Nodal*, *Cerberus1*, *Eomesodermin* and *Foxa2* in E6.5 wildtype and *Axin1^ΔEpi^;Axin2^-/-^*embryos. **(C)** Heatmap showing the scaled gene expression of differentially expressed genes in nascent mesoderm #1, nascent mesoderm #2 and posterior primitive streak cells. The cell type and genotype annotation for each cell (row) is indicated by the colored bars to the left of the heatmap.

Notably, both E6.5 and E7.5 se-bulk RNAseq analysis revealed that targets of the NODAL-signaling pathway such as *Nodal1*, *Cerberus1* and *Lhx1-* were downregulated in the transcriptomes of *Axin1^ΔEpi^;Axin2^-/-^*embryos (**Figure S4D-E** and **Table S2,S4)**. Furthermore, *Lefty1* was downregulated at E6.5, whereas *Lefty2* and *Foxh1*,(Meno et al., 1999; Saijoh et al., 2000) were downregulated in the E7.5 *Axin1^ΔEpi^;Axin2^-/-^* se-bulk transcriptome (**Figure S4D-E** and **Table S2,S4)**. As was observed in **Figure 2B** and **2F**, the anterior primitive streak cell cluster was almost absent in E6.5 *Axin1^ΔEpi^;Axin2^-/-^*mutant compared to control embryos; suggesting that the down regulation of NODAL-signaling target genes found in the se-bulk RNA analysis of *Axin1^ΔEpi^;Axin2^-/-^* embryos was driven by a shift in cell populations (**Figure 4B**). Taken together these data suggest that changes in gene expression occurring in the mutant cell populations shift cells away from an anterior primitive streak cell identity in both *Axin1^ΔEpi^;Axin2^-/-^*and *Smad2* mutant embryos.

Control and mutant embryos contained three overlapping clusters: nascent mesoderm #1, nascent mesoderm #2 and posterior primitive streak cells (**Figure 2E-F** and **Figure S2A**). To explore the impact of constitutive WNT signaling on gene expression in these populations we identified genes that were differentially expressed in control versus *Axin1^ΔEpi^;Axin2^-/-^*cells (**Figure 4C**, control = blue, *Axin1^ΔEpi^;Axin2^-/-^* = red). These analyses revealed that ectopic WNT signaling drove increased expression of the transcription factors *Lmo2* and *Hoxb1* (**Figure 4C**, pink arrowheads) in nascent mesoderm populations #1 and #2. *Lmo2* is an essential regulator of yolk sac hematopoiesis (Warren et al., 1994); its expression is regulated by HOX proteins in embryonic mesoderm at later stages of embryogenesis (Calero-Nieto et al., 2013). Strikingly, the transcription factor *Mixl1* was also overexpressed in *Axin1^ΔEpi^;Axin2^-/-^* nascent mesoderm (**Figure 4C**, pink arrowheads). A previous study demonstrated that overexpression of MIXL1 in ES cell-derived embryoid bodies, accelerates mesoderm formation and increases output of hematoendothelial progenitors (Willey et al., 2006). *Axin1^ΔEpi^;Axin2^-/-^* posterior primitive streak cells showed increased expression of caudal type homeobox transcription factors *Cdx1/2/4* and downregulation of the pluripotency-associated transcription factor *Pou5f1* (**Figure 4C**, pink arrowheads). CDX4, when overexpressed, alters *Hox* gene expression in zebrafish and induces blood formation in mouse ES cultures (Davidson et al., 2003). se-bulk RNAseq analysis also revealed that *Lmo2*, *Mixl1* and *Cdx1/2/4* were upregulated in E7.5 *Axin1^ΔEpi^;Axin2^-/-^* embryos (**Table S3**). These findings align with the increase in hematoendothelial progenitors in the *Axin1^ΔEpi^;Axin2^-/-^* mutants. Furthermore, consistent with loss of the *Axin1* and *Axin2* inhibitors in nascent mesodermal cells, both E6.5 scRNAseq and se-bulk transcriptome analysis showed increased expression of *Dkk1, a* negative regulator of WNT signaling (Niida et al., 2004) in the mutants (**Figure 4C**, pink arrowhead and **Table S2**). Taken together these data suggest that *in vivo*, constitutively active WNT signaling blocks the generation of anterior primitive streak and dysregulates gene expression programs in nascent mesoderm cells towards ExM/posterior LPM markers.

### In the absence of the BMP component of the WNT-BMP regulatory circuit, excess WNT signals drive epiblast differentiation into anterior primitive streak

We next investigated the role of increased BMP signaling in driving the ExM/posterior LPM gene expression program in the *Axin1^ΔEpi^;Axin2^-/-^* mutant. We hypothesized that perturbation of the BMP-WNT regulatory circuit in *Axin1^ΔEpi^;Axin2^-/-^* mutants generates changes in gene expression that result in the repression of signals, such as NODAL/pSMAD2/3, required to produce anterior primitive streak progenitors. To test this hypothesis, we used a conditional *Bmpr1a* allele to reduce levels of BMP signaling in the epiblast of *Axin1^ΔEpi^;Axin2^-/-^* embryos (**Figure 5A**). The resulting triple mutant-*Axin1^ΔEpi^;Axin2^-/-^;Bmpr1a^ΔEpi^*-embryos reached E7.75 and displayed a distinct morphology from that of *Axin1^ΔEpi^;Axin2^-/-^*embryos at E7.75 (**Figure 5B**). Moreover, in contrast to *Axin1^ΔEpi^;Axin2^-/-^*embryos; the *Axin1^ΔEpi^;Axin2^-/-^;Bmpr1a^ΔEpi^* triple mutants differentiated TBX6-expressing cells (**Figure 5C,D**). An analysis of the TCF/Lef:H2B-GFP WNT reporter and IF for pSMAD1/5/9 expression in *Axin1^ΔEpi^;Axin2^-/-^;Bmpr1a^ΔEpi^* triple mutants demonstrated active WNT signaling throughout the embryonic region and a near complete absence of BMP pathway activity (**Figure 5D-E)**. The *Axin1^ΔEpi^;Axin2^-/-^;Bmpr1a^ΔEpi^* triple mutant also contained reduced numbers of KDR-positive cells (**Figure 5D-E**). Thus, decreasing the levels of BMP signaling in the epiblast of *Axin1^ΔEpi^; Axin2^-/-^*embryos enabled the differentiation of paraxial mesoderm. Taken together our findings show that removal of the BMP component of the regulatory circuitry, leads to excess WNT signaling that drives epiblast differentiation towards anterior primitive streak cell derivatives.

**Figure 5.**
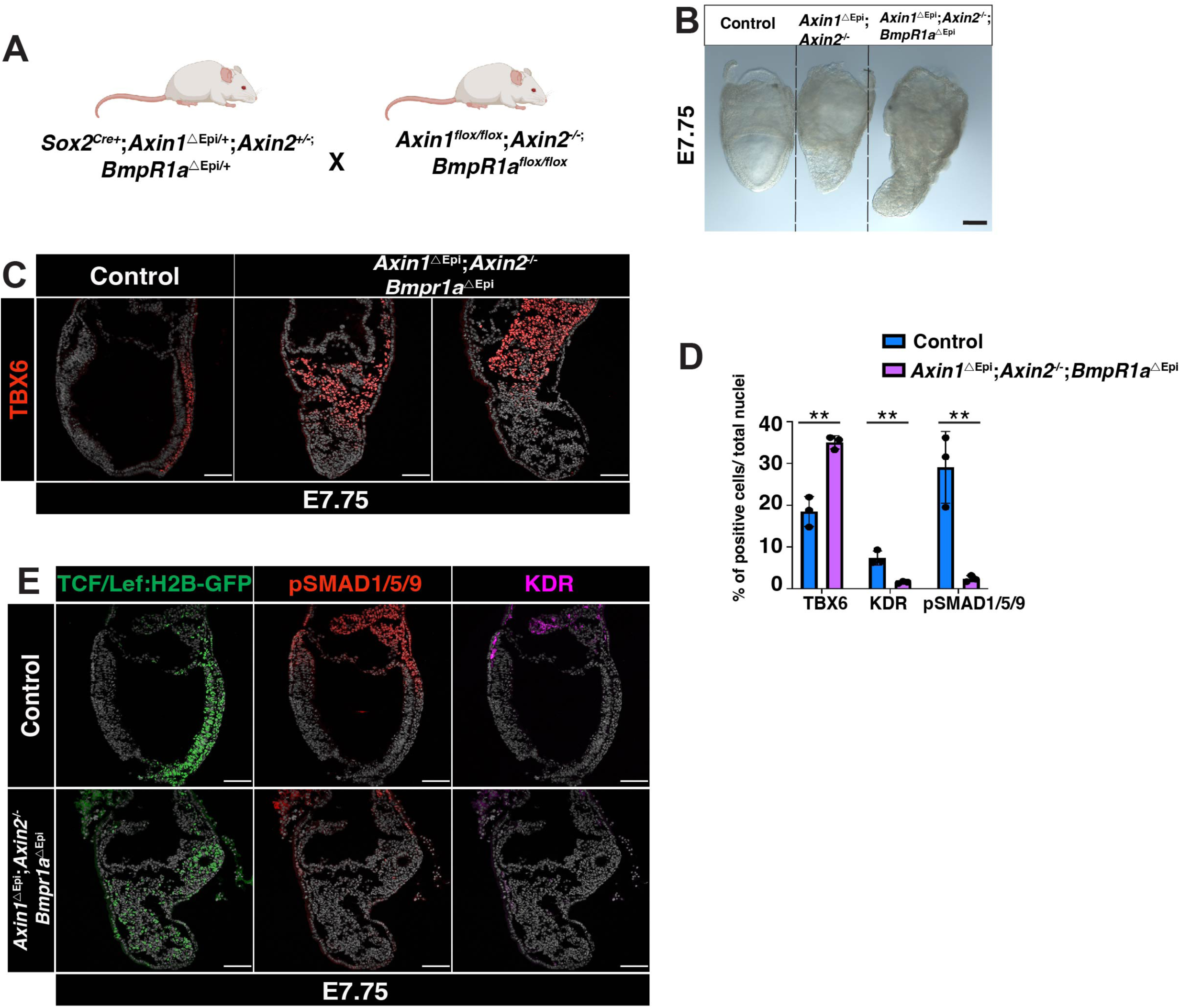
Excess of WNT signals differentiate the epiblast to paraxial mesoderm *in vivo* when BMPR1a is deleted. (**A**) Cartoon describing the mice mating strategy to generate the triple mutants. “Created with BioRender.com”. (**B**) Bright field images of E7.75 wild-type, *Axin1^ΔEpi;^Axin2^-/-^* and *Axin1^ΔEpi^;Axin2^-/-^;Bmpr1a^ΔEpi^* triple mutants. Scale bar is 100μm. (**C**) Immunostainings for TBX6 on E7.75 wild-type and *Axin1^ΔEpi^;Axin2^-/-^;Bmpr1a^ΔEpi^* triple mutants. (**D**) Quantitation of TBX6, KDR and pSMAD1/5/9-positive cells in wild-type and *Axin1^ΔEpi^;Axin2^-/-^ ;Bmpr1a^ΔEpi^*triple mutants embryos. Bars represent the mean± SEM. Statistically significant differences were determined by t-test. **p<0.005. (**E**) IFs for pSMAD1/5/9, GFP (TCF/Lef:H2B-GFP) and KDR in wild-type and *Axin1^ΔEpi^;Axin2^-/-^;Bmpr1a^ΔEpi^* E7.75 longitudinal sections. Scale bars on IFs are 50μm.

### Differentiation of anterior and posterior primitive streak derivatives occurs *in vitro* upon activation of WNT-signaling

To further investigate the kinetics of WNT activation in specifying mesoderm, we derived *Axin1^fus/fus^;Axin*2^-/-^ double mutant Embryonic Stem Cells (ESCs) from blastocyst stage embryos and transitioned these, along with *Smad2^-/-^* ESCs, towards an Epiblast-Like Cell (EpiLC) state. Whereas ESCs reside in a naïve pluripotent state, EpiLC recapitulate a later formative pluripotent transcriptional state, approximating the post-implantation epiblast prior to the onset of gastrulation (Hayashi et al., 2011). When cultured in either 2i or serum plus LIF, *Axin1^fus/fus^;Axin*2^-/-^ ESCs expressed the markers characteristic of naïve state pluripotency (**Figure S5A**). Following 24 hours in FGF2 + Activin A, neither wild-type nor *Axin1^fus/fus^;Axin*2^-/-^ EpiLCs expressed detectable levels of T protein; based on IF staining (**Figure S5B)**. However, by 48 hours, *Axin1^fus/fus^;Axin*2^-/-^, but not wild-type, EpiLCs initiated their transition toward mesoderm differentiation (**Figure S5B**). Following 48 hours of differentiation, the *Axin1^fus/fus^;Axin2^-/-^* mutant EpiLCs initiated a premature EMT program, characterized by the expression of T and CDH2, and the downregulation of CDH1. By contrast, wild-type and *Smad2* null EpiLCs never expressed T or CDH2 under the same conditions (**Figure S5C**). Notably after 72 hours *Axin1^fus/fus^;Axin*2^-/-^,EpiLC expressed even higher levels of aberrant T and CDH2 expression, whereas wild-type EpiLCs never underwent a gastrulation EMT when cultured under EpiLC conditions (**Figure 6A-B**).

**Figure 6.**
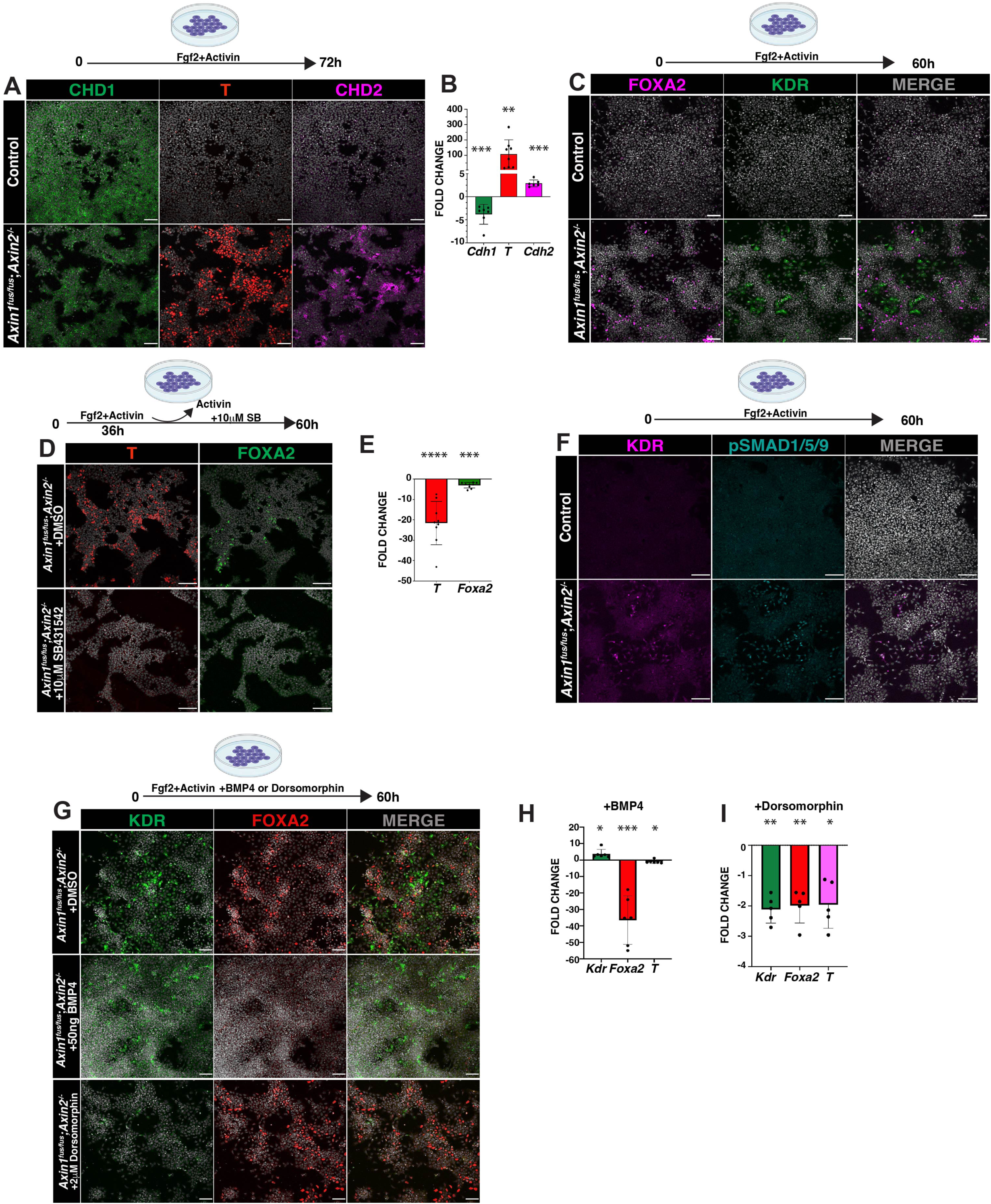
Differentiation of anterior and posterior mesoderm progenitors occurs *in vitro* upon activation of WNT signaling. (**A**) IF on wild-type and *Axin1;Axin2* EpiLC mutant cells for CDH1, T and CDH2 after differentiating for 72 hours. (**B**) Graph showing expression of *T*, *Cdh1*, and *Cdh2* in *Axin1;Axin2* mutant EpiLC compared to wild-type detected by qPCR. (**C**) IF on wild-type and *Axin1;Axin2* mutant EpiLC for FOXA2 and KDR after differentiating for 60 hours. IFs for T and FOXA2 on *Axin1;Axin2* mutant EpiLC plus DMSO and *Axin1;Axin2* mutant EpiLC treated with Nodal-receptor inhibitor during 24 hours. (**E**) Graph showing levels of *T* and *Foxa2* expression detected by qPCR in *Axin1;Axin2* mutant EpiLC after Nodal receptor inhibitor-treatment relative to control. (**F**) IF on wild-type and *Axin1;Axin2* mutant EpiLC for KDR and pSMAD1/5/9 after 60 hours of differentiation. (**G**) IF on *Axin1;Axin*2 mutant cells for KDR and FOXA2 after grown for 60 hours with either BMP4 or Dorsomorphin. Scale bars are 50μm. (H) Graph showing levels of *Kdr*, *Foxa2* and *T* in *Axin1;Axin2* mutant cells after treatment with either BMP4 or Dorsomorphin determined by qPCR. Statistically significant differences were determined by t-test. *p<0.03, **p<0.009, ***p<0.001,****p<0.0001. All the cartoons on top of the panels describe the experimental strategy and were “Created with BioRender.com”.

Reflective of the necessary role that NODAL/pSMAD2/3 plays in the generation of all mesodermal lineages *in vivo* in the mouse embryo, the protocol for EpiLC differentiation *in vitro* requires the addition of exogenous Activin A, a commercial NODAL agonist. We hypothesized that *Axin1^fus/fus^;Axin2^-/-^*mutant EpiLCs would be capable of differentiating towards mesoderm derived from the anterior streak. To test this hypothesis we assayed FOXA2 expression and observed that while *Axin1^fus/fus^;Axin2^-/-^*mutant EpiLCs differentiate axial mesoderm, wild-type EpiLCs did not express FOXA2 (**Figure 6C**).

Considering that *Smad2* mutant embryos fail to differentiate mesodermal cells of the anterior streak, and that canonical WNT signaling induces *Nodal* via an identified proximal epiblast enhancer (PEE) (Ben-Haim et al., 2006) we asked whether the specification of FOXA2 in *Axin1^fus/fus^;Axin2^-/-^*EpiLCs requires the NODAL/SMAD2/3 signaling pathway, possibly acting downstream of WNT. We cultured *Axin1;Axin*2 mutant EpiLCs for 36 hours to allow them to differentiate to mesoderm; after that time interval we withdrew Activin and added an ALK-inhibitor (SB431542) for 24 hours (a total of 60 hours) and then we analyzed FOXA2 expression by IF. We found an ∼30 fold reduction of T and an ∼3 fold reduction of FOXA2 (**Figure 6D,E**), indicating that *Axin1;Axin*2 mutant EpiLCs require a functional NODAL/SMAD2/3 pathway to differentiate ectopic axial mesoderm. Since *Axin1^ΔEpi^;Axin2^-/-^* embryos aberrantly differentiate towards ExM/posterior LPM, we investigated the ability of *Axin1^fus/fus^;Axin2^-/-^* EpiLCs to form mesoderm in these lineages by monitoring KDR expression. After 60 hours we detected KDR-positive cells in *Axin1;Axin2* mutant EpiLCs that express FOXA2 at the same time (**Figure 6C**). These findings suggest that overactivation of WNT promotes EpiLC differentiation towards both anterior and posterior mesodermal lineages.

Observing KDR+ cells, prompted us to assess the status of the BMP signaling pathway in the mutant EpiLCs. We observed pSMAD1/5/9-positive cells only at the edge of the *Axin1;Axin*2 mutant EpiLC colonies after 60 hours, whereas most of the remaining mutant EpiLCs did not activate the BMP/pSMAD1/5/9 pathway (**Figure 6F**). As predicted, the pSMAD1/5/9-positive cells also expressed KDR, a marker of ExM identity (**Figure 6F**). By contrast, wild-type cells did not activate the BMP/pSMAD1/5/9 pathway, nor did they differentiate towards ExM (**Figure 6F**).

We note that ExM differentiation, detected by KDR expression, occurred at the edges of the EpiLC colonies or in cells having undergone EMT, whereas axial EM differentiation denoted by FOXA2+ expression occurred in the center of the EpiLC colony (**Figure 6C**). The majority of the *Axin1;Axin*2 mutant EpiLCs expressed a mesoderm marker labeled for either KDR or FOXA2 but not both; we observed only a few cells expressing both markers.

*Axin1^ΔEpi^;Axin2^-/-^* mutant embryos fail to differentiate derivatives of the anterior primitive streak, while *Axin1;Axin*2 mutant EpiLCs expressed FOXA2. Human ESCs grown in high-density colonies have been shown to localize their BMP receptors on their lateral cell adjacent surfaces within the center of the colony, while apically localizing of the receptors on the outer edges. This differential localization insulates cells at the center from apically applied ligands (Etoc et al., 2016). To reconcile our apparently contradictory observations *in vivo* in embryos versus *in vitro* in EpiLCs, we hypothesized that similar to human ESCs, mouse EpiLC are more sensitive to BMP signals at the edge of the colony. At limiting concentrations of BMP ligand cells on the edges of EpiLC colonies would differentiate towards ExM, while cells at the center would differentiate towards FOXA2, unable to access levels of ligand needed to activate the BMP signaling pathway. To test this idea, we maintained *Axin1;Axin*2 mutant EpiLCs in the presence of BMP4 for 60 hours and then evaluated their differentiation towards both posterior and anterior mesoderm lineages. We observed increased posterior mesoderm differentiation in *Axin1;Axin*2 mutant EpiLCs exposed to higher concentrations of BMP4, demonstrated by the ∼4 fold increase in KDR expression (**Figure 6H** 3.7 average fold). Under these conditions, we detected KDR-positive cells not only at the edges of the *Axin1;Axin*2 mutant EpiLCs colonies, but also in the center (**Figure 6G**). As predicted, differentiation towards FOXA+ cells in mutant EpiLC was dramatically reduced in the presence of high levels of BMP (∼36.5 average fold decrease) (**Figure 6G-H**). EpiLC were cultured in the presence of Dorsomorphin, a BMP-receptor inhibitor, to test whether we could modulate *Axin1^fus/fus^;Axin*2^-/-^ EpiLC differentiation towards axial mesoderm by inhibiting the BMP/pSMAD1/5/9 pathway. We observed that Dorsomorphin reduced *Kdr* expression in mutant EpiLC by about 2 fold (**Figure 6I**). However, *Foxa2* expression did not change by IF (**Figure 6G**), but we detected a decrease in *Foxa2* expression by qRT-PCR, likely due to a general decrease in mesoderm specification, corroborated by a decrease in T-expression (**Figure 6I**).

The findings from our EpiLC experiments suggest that overactivation of WNT signaling is sufficient to induce mesoderm differentiation. The ectopic differentiation of either posterior or anterior mesodermal lineages depends on mesodermal progenitors attaining a threshold level of either WNT+BMP/pSMAD1/5/9 or WNT+NODAL/pSMAD2/3 signaling respectively.

## DISCUSSION

A long-term goal of our research is to uncover the molecular and cellular mechanisms that coordinate the emergence of mesodermal lineages from epiblast cells that reside in and move through the proximal, posterior primitive streak early during gastrulation. In the present study we asked whether specification of distinct mesodermal lineages involves an interplay between NODAL/BMP pathway activities and dose/stage-dependent WNT signaling. We achieved deregulation of WNT in mouse embryos and EpiLCs using loss of function mutations in *Axin1* and *Axin2,* two negative regulators of WNT signaling. Applying bulk and scRNAseq on control and *Axin1/2* mutant embryos, this study has identified a previously undescribed role for canonical WNT signaling in the specification of ExM, posterior LPM and intermediate mesoderm from pluripotent epiblast during gastrulation. This role includes two layers of regulation: first, WNT initiates the differentiation of primitive streak cells into mesoderm progenitors; second, WNT amplifies and cooperates with BMP/pSMAD1/5/9 and NODAL/pSMAD2/3 to propel differentiating mesoderm progenitors into, respectively, posterior plus middle streak derivatives and anterior streak derivatives.

Importantly our findings will contribute to ongoing discussions on the hierarchical versus combinatorial nature of the regulatory networks driving lineage specification during gastrulation. Recent studies using transcriptomic approaches have challenged the notion of a binary cell fate decision giving rise in the epiblast to presumptive progenitors of either ExM or embryonic mesoderm (Parameswaran and Tam, 1995). Instead they suggest that the extraembryonic, lateral, intermediate, paraxial and axial subtypes of mesoderm arise from independent progenitors (Mittnenzweig et al., 2021; Scialdone et al., 2016). Our observations align with the more recent view of regulatory networks since our analyses did not detect two sharp transitions between ExM or embryonic, hinting at a more complex combinatorial interplay of signaling pathways operating in the epiblast to generate diverse mesodermal cell types.

Here we found that when constitutively active, the WNT pathway interacts with BMP signaling to promote aberrant differentiation of the epiblast and nascent mesoderm into ExM, posterior LPM and intermediate mesoderm. Specification of the LPM and intermediate mesoderm lineages occur shortly after their epiblast progenitors leave the primitive streak during gastrulation (Wilson and Beddington, 1996). However, the limited molecular signatures associated with LPM and intermediate cells have precluded defining their precise segregation (Prummel et al., 2020). Importantly the sc-datasets contributing to the discovery of a combined role for WNT/BMP in the specification of posterior LPM and intermediate mesoderm, provide a cohort of genes modulated by both signals. These provide promising candidates to pursue in studies seeking to elucidate the precise temporal specification of these mesoderm lineages during early mouse gastrulation.

An extensive body of research on vertebrate gastrulation has explored the role of feedback loops operating between cells to amplify small differences and generate stable patterns. In mouse, studies characterizing the phenotypes displayed by embryos carrying mutations in individual components of the BMP, WNT and NODAL signaling pathways produced a working model positing that BMP-WNT and WNT-Nodal regulatory feedback loops achieve the posterior position of the primitive streak (Ben-Haim et al., 2006). Here we report that a BMP-WNT positive regulatory feedback loop operates to promote ExM, posterior LPM and intermediate mesoderm differentiation. We show that this WNT-BMP regulatory circuit is amplified in *Axin1^ΔEpi^;Axin2^-/-^* and *Smad2^-/-^*mutant embryos, and that when active in either epiblast or primitive streak cells, it consistently results in ectopic ExM differentiation. Confirming the central role of BMP in this regulatory circuit, we find that even in the presence of constitutively active WNT signaling, embryos lacking *Bmpr1a* -as in the *Axin1^ΔEpi^;Axin2^-/-^;Bmpr1a^ΔEpi^* triple mutant-are not able to ectopically differentiate ExM. In our *in vivo* studies, we localized cells actively transducing BMP signals to the most proximal region of the primitive streak in wild-type embryos. Derivatives of the anterior streak express the BMP antagonists Noggin and Chordin (McMahon et al., 1998), suggesting that they act during normal development to prevent anterior cells from responding to BMP, thereby limiting BMP signaling activity to the proximal epiblast. Here we show that constitutively active WNT signaling increases the expression of different BMP ligands in the epiblast of both *Axin1^ΔEpi^;Axin2^-/-^*and *Smad2^-/-^* embryos; the increased availability of ligand likely contributed to the expanded region of pSMAD1/5/9-BMP activity.

The NODAL/pSMAD2/3 pathway plays a fundamental role in directing the temporal and spatial pattern of specification and differentiation of the multiple mesoderm and endoderm lineages that arise from the primitive streak during gastrulation (Dunn et al., 2004; Vincent et al., 2003). The NODAL-signaling pathway is severely downregulated in global loss of function *Smad2^-/-^*mutant embryos and the majority of epiblast cells abnormally differentiate into ExM, a phenotype resembling that observed in *Axin1^ΔEpi^;Axin2^-/-^*mutant embryos. The similarly strong downregulation of the NODAL pathway detected by se-bulk RNAseq analyses of *Axin1^ΔEpi^;Axin2^-/-^*mutants results from the lack of anterior streak progenitors, as revealed by the scSeq analysis. Genetic studies *in vivo* have allowed us to investigate the consequences of perturbed WNT and NODAL signaling in the context of normal and reduced BMP pathway activity. Notably, our analysis of the *Axin1^ΔEpi^;Axin2^-/-^;Bmpr1a^ΔEpi^* triple mutant revealed that inhibition of BMP signaling restores paraxial (TBX6-positive) mesoderm, normally absent in *Axin1^ΔEpi^;Axin2^-/-^* mutant embryos.

To elucidate the interplay between the WNT, BMP and NODAL pathways in posterior streak derivatives (KDR+) versus anterior streak differentiation (FOXA+), we employed the *in vitro* EpiLC model. Our results suggest that WNT signals must reach a threshold of activation (24 hours) before *Axin1;Axin2* mutant EpiLCs initiate expression of *T*, a marker of mesodermal progenitors. Finding that generation of differentiated mesoderm requires an additional ∼26 hours of *Axin1;Axin2* EpiLC culture, also intimates that ectopic differentiation of either KDR+ or FOXA2+ cells depends on mesodermal progenitors attaining a threshold level of either WNT+BMP/pSMAD1/5/9 or WNT+NODAL/pSMAD2/3 signaling. We showed that pharmacological inhibition of the NODAL-signaling pathway decreases ectopic differentiation of FOXA2+ population; conversely, inhibition of BMP signaling blocks the differentiation of *Axin1;Axin2* mutant EpiLCs towards KDR+ cells, confirming the interdependence of the WNT+NODAL/pSMAD2/3 and WNT+BMP/pSMAD1/5/9 pathways in anterior and posterior mesoderm specification respectively. Our *in vitro* studies also shed light on the combinatorial interpretation of multiple signals by epiblast cells to generate the different mesodermal lineages. The EpiLC system might help to investigate how WNT signaling impacts complex and rich cell behaviors (ranging from cell adhesion, cell shape changes, ligand concentration) that alter gene expression patterns and propel a mesodermal progenitor into a posterior versus anterior lineage.

During gastrulation, cells experience simultaneous signaling through multiple pathways with dynamically changing activity. This work refines our mechanistic understanding of how the WNT-NODAL-BMP pathways interact *in vivo* at the primitive streak to modulate differential cell fate allocation. The work also has implications for mouse and human stem cell-based embryo models (e.g. 3D gastruloids) which rely on a pulse of WNT activity to break radial symmetry and induce axial elongation (Van Den Brink et al., 2014).

## METHODS

### Mouse strains

*Axin1* null mutants were generated from a knockout-first allele (*Axin1^tm1a(KOMP)Wtsi^*) derived from a mouse embryonic stem cell (ESC) line obtained from The International Mouse Phenotyping Consortium (IMPC, https://www.mousephenotype.org/). ESCs carrying the *Axin1^tm1a(KOMP)Wtsi^* allele were injected into embryos to generate germline transmitting chimeras by the MSKCC Mouse Genetics Core Facility. To generate the null allele, the conditional allele of *Axin1* was crossed to *CAG-Cre* transgenic animals (Sakai and Miyazaki, 1997) (Jackson Laboratory). The *Axin1^fus^* (Axin1-fused) allele was previously described (Zeng et al., 1997b). *Axin2^tm1Wbm^*(lacZ knock-in) (Lustig et al., 2001b) was obtained from Frank Constantini (Columbia University, New York). Mice carrying the *Axin1* floxed allele (*Axin1^flox/flox^*) were crossed with *Sox2-Cre* (Hayashi et al., 2002); *Axin2*^+/-^ mice to generate males carrying the desired allele combination (*Sox2-Cre^Tg/^*^+;^ *Axin1^ΔEpi/+^;Axin2^+/-^*). We obtained a *Smad2* conditional allele (*Smad2^tm1.1Epb^/J*)(Ju et al., 2006) from Jackson Laboratory and generated a *Smad2* null allele (*Smad2^-/-^*) by crossing it to *CAG-Cre* transgenic mice. *Bmpr1a* floxed allele (Mishina et al., 1995) was mated to *Sox2-Cre^Tg/^*^+;^ *Axin1^ΔEpi/+^;Axin2^+/-^*to generate triple carriers: *Sox2-Cre^Tg/^*^+;^ *Axin1^ΔEpi/+^;Axin2^+/-^; Bmpr1a^ΔEpi/+^. Bmpr1a^flox/flox^* mice were crossed to *Axin1^flox/flox^;Axin2^-/-^*to generate *Bmpr1a^flox/flox^;Axin1^flox/flox^;Axin2^-/-^*mice. Either *Sox2-Cre^Tg/^*^+^;*Axin1^ΔEpi/+^;Axin2^+/-^*or Smad2^+/-^ mice were mated to TCF-LEF-H2B-GFP mice to generate *Sox2-Cre^Tg/^*^+;^ *Axin1^ΔEpi/+^;Axin2^+/^;* TCF-LEF-H2B-GFP^+/-^ or Smad2^+/-^; TCF-LEF-H2B-GFP^+/-^. 8-16-week-old female mice were bred to generate embryos recovered at E5.5-E7.75. All mice were maintained in a mixed genetic background. Mice were housed and bred under standard conditions in accordance with IACUC guidelines.

Primers for genotyping the Axin1-null allele are Forward 5’-TGGGATTAAAGGCAGACACC-3’ and Reverse 5’-TCCTGCCTAACTTACACTCTCCTGC-3’ and to genotype the Axin1 floxed allele are: Forward 5’-TGGGATTAAAGGCAGACACC-3’ and Reverse 5’-TGTATGCATGCAGGCAATGACAGG-3’. Primers to genotype Axin2 null allele are: Axin2-9391 AAGCTGCGTCGGATACTTGCGA; Axin2-9392 AGTCCATCTTCATTCCGCCTAGC and Axin2-9393 TGGTAATGCTGCAGTGGCTTG. Primer to genotype the Smad2 null allele are: Forward TTCCATCATCCTTCATGCAAT and Reverse: GACCAAGGCGAAAGGAAACT. Primers to genotype BmpR1a floxed allele are: Bmpr1a-Fx1-GGTTTGGATCTTAACCTTAGG; Bmpr1a-Fx2 GCAGCTGCTGCTGCAGCCTCC and Bmpr1a-Fx3 TGGCTACAATTTGTCTCATGC

### Immunostaining on cryosections

Immunofluorescent staining of embryo sections and whole mount embryos was performed according to standard protocols. Briefly: embryos were fixed for 1 hour in 4% PFA on ice followed by 4 washes with PBS, 10 minutes each. For sectioning, embryos were incubated in 30% sucrose in PBS on ice until they sank. Embryos were embedded in OCT (Tissue-Tek) frozen and cryosectioned at 10μm-12μm. Immunostaining was performed in blocking buffer: PBS containing 4% heat-inactivated donkey serum (Gemini, Bio-products) and 0.1% Triton X-100. The following primary antibodies were incubated overnight in blocking buffer: CDH1 (1:300 Sigma-Aldrich), CDH2 (1:300, Cell Signaling Technology), T (1:400, Cell Signaling), KDR (1:200, BD Pharmigen), CDX2 (1:400, Cell Signaling), FOXF1 (1:300, R&D Systems), TBX6 (1:300, R&D Systems), pSMAD1/5/9 (1:100 Cell Signaling), FOXA2 1:500 (Abcam), GFP (1:500, R&D Systems), SOX2 (1:400 Cell Signaling), POUF5F1 (1:500, SantaCruz), NANOG (1:300 BD Biosciences). Fluorescent-secondary antibodies (Invitrogen) were diluted at 1:400 in blocking buffer and incubated for 1 hour at RT (room temperature). The mouse embryo sections were mounted in ProLong Gold (ThermoFisher) and imaged either using a Leica-upright SP5 laser point-scanning microscope or a Zeiss LSM 880 laser-scanning confocal microscope. Confocal images were reconstructed and analyzed by using Volocity software package (PerkinElmer) and Fiji. Raw image data was analyzed using Fiji (Rasband, W.S, Image J, U.S. National Institutes of Health, Bethesda, Maryland, USA.)

### mRNA *in situ* hybridization

*In situ* hybridization was performed according to protocols previously described (Eggenschwiler and Anderson, 2000). Briefly: the embryos were fixed overnight in 4% PFA. Next day, the embryos were washed in PBS with 0.1% Tween, dehydrated in graduated methanol series. The following day the embryos were hydrated in graduated methanol series, incubated in 10μg/ml of PK/PBS solution for 3-7 minutes and hybridized in hybridization solution plus RNA probe at 70°C overnight. A series of washes in 2XSSC and MAB solutions (100mM Maleic acid, 150mM NaCl in PBS) was performed next day. The embryos were incubated in 1:1000 anti-DIG antibody (Roche) in 10% Goat-Serum; 0.1% Tween-20 in PBS overnight and then washed extensively in PBT (0.1% BSA; 0.1% Tween-20 in PBS). The embryos were incubated in BM purple solution (Roche) until the development of purple color.

### Quantitation of T, CDH2, TBX6, CDX2, FOXF1, KDR, pSMAD1/5/9 and TCF/Lef:H2B-GFP-positive cells

Fiji (Schindelin et al., 2012) was used to manually count the number of nuclei expressing the different markers in a single Z-plane per embryo from a stack of the wild-type and the mutant sections. The percentage of positive cells per each marker was determined from the number of positive cells located in a single z-plane divided by the total number of DAPI+ nuclei in that Z-plane. We performed the same measurements in 4 alternate cryo-sections from each wild-type and mutant embryos collected and average the percentage per embryo. The mutant and the wild-type embryo sections were stained at the same time. At least three different wild-type and mutant embryos were analyzed.

### Quantitation of T, SOX2, NANOG and POU5F1 positive cells

Fiji (Schindelin et al., 2012) was used to manually count the number of nuclei expressing the different markers in a single Z-plane per embryo from a Z-stack of the wild-type and the mutant sections. The percentage of positive cells per each marker was determined from the number of positive cells located in a single z-plane divided by the total number of DAPI+ nuclei in that Z-plane. We performed the same measurements in the whole Z-stack from either E5.5 or E5.75 wild-type and mutant embryos collected and average the percentage per embryo. The mutant and the wild-type embryos were stained at the same time. At least three different wild-type and mutant embryos were analyzed.

### Bulk RNA-sequencing and data analysis

(4) E6.5 wild-type and (4) E6.5 *Axin1^ΔEpi^;Axin2^-/-^* mutant embryos, and (5) E7.5 wild-type and (5) E7.5 *Axin1^ΔEpi^;Axin2^-/-^* mutant embryos were dissected in cold PBS, and individually collected in 500µl of TRIzol reagent (Invitrogen). RNA-sequencing libraries were prepared and processed by the IGO Core Facility (MSKCC). SMARTer total RNA-seq kit was used to generate RNA-seq libraries for Illumina sequencing. A sequencing depth of between 30-40 million paired end reads per sample were analyzed.

Briefly, output data (FASTQ files) was mapped to the mouse genome using STAR aligner (Dobin et al., 2012). A 2 pass mapping method outlined in (Engström et al., 2013). After mapping we post processed the output SAM files using the PICARD tools to: add read groups, AddOrReplaceReadGroups which in additional sorts the file and coverts it to the compressed BAM format. We then computed the expression count matrix from the mapped reads using HTSeq (www-huber.embl.de/users/anders/HTSeq) and one of several possible gene model databases. The raw count matrix generated by HTSeq are then be processed using the R/Bioconductor package DESeq (www-huber.embl.de/users/anders/DESeq) which is used to both normalize the full dataset and analyze differential expression between sample groups.

The data was clustered in several ways using the normalized counts of all genes that a total of 10 counts when summed across all samples.

Hierarchical cluster with the correlation metric (D_{ij} = 1 - cor(X_i,X_j)) with the Pearson correlation on the normalized log2 expression values.

Multidimensional scaling

Principal component analysis

Heatmaps are generated using the heatmap.2 function from the gplots R package. For the Heatmaps the top *100* differentially expressed genes are used. The data plot was the mean centered normalized log2 expression of the top 100 significant genes. For mouse datasets we run a gene set analysis using GSEA with gene sets from the Broad <u>mSigDb</u>. If a sample group has fewer than three samples, we run GSEAPreranked using log2 fold changes from DESeq.

### Single-cell (sc)RNA-sequencing

(15) E6.5 control and (10) E6.5 *Axin1^ΔEpi^;Axin2^-/-^* double mutant embryos were single cell dissociated to generate the sc-RNA libraries. Control and mutant embryos were dissected at the same time. Control embryos were obtained by crossing FVB females (Jackson) to the same males used to obtain the *Axin1^ΔEpi^;Axin2^-/-^* double mutant embryos (Sox2-Cre^+^; *Axin1^ΔEpi/+^;Axin2^+/-^*). Mutant embryos were identified by distinctive morphology and dissected in DMEM-F12 + 5% Newborn Calf Serum (NCS) (Gibco). Individual embryos were washed three times in 40µl drops of DMEM-F12 media. Thereafter, batches of 4 embryos were digested in 40µl of a solution of 2:1 0.25% Trypsin (Gibco): Accutase (StemCell Technologies) at 37C for 15 minutes. After partial digestion, 40µl of (DMEM-F12 + 20% NCS + 4mM EDTA) solution was added to each batch of embryos which were mechanically dissociated by hand pipetting, followed by mouth pipetting with pulled glass capillaries of different diameters to obtain a single cell suspension. The cell suspension was collected in DMEM-F12 + 10% NCS and filtered through a FlowMi cell strainer (4mM Millipore, Sigma). Cells were centrifugated at 2,200 rpm for 4 minutes and washed twice with DMEM-F12 + 10% NCS. Cells were counted and diluted in DMEM-F12 + 10% NCS. Cells were loaded on a Chromium Controller microfluidic chip for encapsulation with a target number of 10,000 cells for the generation of generate single cell 3’ RNA-seq libraries. Libraries were generated following the manufacturer’s instructions (10x Genomics Chromium Single Cell 3’ Reagent Kit User Guide v2 Chemistry), as previously (Nowotschin et al., 2019).

### Gene ontology analyses

Data from bulk-RNA sequencing was analyzed using the Gene Ontology website (http://geneontology.org). A list of the most representative terms with a 3-fold change or higher either up-regulated or down-regulated was selected to include in the figures.

### Statistics and Graphs

Statistical analysis and graphs were performed using Prism (Graph Pad). Graphs of changes in fold expression from the mouse embryos were plotted using the data from the se-bulk-RNA sequencing. T-test statistical analysis was performed to obtain the statistical significance.

### Mapping scRNA sequencing data & pre-processing

Raw sequencing files were mapped to the mm10 reference genome (10x Genomics refdata-cellranger-mm10-1.2.0) using CellRanger count (v7.0.1) with chemistry=SC3Pv3. High quality cells were retained using the following thresholds: log10(number of reads) > 3.5 & < 5, number of genes > 1.5e3 & < 1e4, percentage mitochondrial RNA < 5%, percentage ribosomal RNA < 35%. Doublet calling was performed with the scds ((Bais and Kostka, 2020) v1.14.0) package using the hybrid approach and doublet score > 1 as threshold. A final step of quality control was added to remove cells with percentage mitochondrial RNA >0.8% & < 5%. Count normalization was performed using NormalizeCounts in Seurat v4 with default settings. A total of 10,250 cells passed quality control.

### Integration and UMAP generation for the Control and *Axin1^ΔEpi^;Axin2^-/-^* datasets

The control and Axin-DKO scRNA-seq datasets were integrated using the Reciprocal Principal Component Analysis (rPCA) pipeline in Seurat v4, with parameters set to nfeatures = 2000, k.anchor = 5, and k.weight = 100. The integrated data underwent preprocessing with ScaleData, followed by dimensionality reduction using RunPCA and RunUMAP (Uniform Manifold Approximation and Projection) with ndims = 1:50. Cluster identification was performed using FindNeighbors (ndims = 1:20) and FindClusters (resolution = 1).

### Integration of the Axin datasets with the Pijuan-Sala 2019 gastrulation reference atlas

Using the log normalized counts from a down sampled version (10,000 cells per embryonic stage from E6.5 to E8.5) of a mouse gastrulation atlas from Imaz-Rosshandler et al., 2024 we performed two subsequent rounds of rPCA integration using Seurat v4. In round 1, we performed rPCA integration across different samples (sequencing batches) within each embryonic stage (e.g. E6.5) using 2000 highly variable features identified with FindVariableFeatures and then ran FindIntegrationAnchors (k.anchor = 5, scale = FALSE) and IntegrateData (features.to.integrate = all features (genes), k.weight = 100). In round 2, batch corrected counts matrices from round 1 (9 datasets in total from embryonic stages E6.5 – E8.5) were integrated together again using rPCA integration from Seurat v4. To perform round 2 rPCA integration, features were set to the unique combined list of all *VariableFeatures* identified from each embryonic stage during round 1 and the rPCA parameters (k.weight, k.anchor, scale) in round 2 remained the same as in round 1, described above.

To create a combined integrated Seurat object that contains all cells from the down sampled mouse gastrulation atlas and the axin datasets, query datasets (control and Axin DKO datasets) were integrated with the rPCA batch corrected gastrulation atlas from round 2 (described above), again using rPCA integration (k.anchor = 20 and k.weight = 200, *reference =* gastrulation atlas, scale = FALSE).

### UMAP, Joint Clustering and Label Transfer using the Combined Integrated Mouse Gastrulation Atlas and Axin Datasets

A joint UMAP, containing the query datasets and the down sampled gastrulation atlas, was generated by scaling the combined integrated data, from the multiple rounds of rPCA described above, and performing *RunPCA* and *RunUMAP* (dims = 1:50). Joint clusters across the combined datasets were identified using *FindNeighbors* and *FindClusters* (dims = 1:30, reduction = “pca”, resolution = 2). Embryonic stages and cell type annotations were transferred to query datasets from the gastrulation atlas using *FindTransferAnchors* and *TransferData (dims = 1:30, reference.reduction = “pca”)*. Next, we generated cell type annotations for the joint clusters using both marker gene expression patterns as well as majority cell type labels from the label transfer. Notably, a joint cluster that contained primordial germ cells (PGC) as well as primitive streak cells was reassigned the annotation posterior primitive streak.

### Identification of Marker Genes and Differentially Expressed Genes

For the query datasets alone, marker genes were determined for the joint-clusters using FindAllMarkers (slot = data) from Seurat v4. Differentially expressed genes in the overlapping populations (Nascent mesoderm #1, Nascent mesoderm #2 and Posterior Primitive Streak) in the Control versus *Axin1^ΔEpi^;Axin2^-/-^* were determined using *FindMarkers* and the top up- and down-regulated genes (slice_head (n = 15), avg_logFC > 0.5, p < 0.5) from each population were selected to generate the heatmap shown in figure 4C using *pheatmap* which was given the original normalized scaled gene expression data of these cell populations from the Control and *Axin1^ΔEpi^;Axin2^-/-^* datasets.

### EpiLC conversion

Embryonic stem cells (ESCs) were derived from E3.5 wild-type and *Axin1^fus/fus^*;*Axin2^-/-^*blastocysts as previously described (Czechanski et al., 2014). ESC lines were maintained in standard serum/LIF ESC medium (Dulbecco’s Eagle’s medium DMEM) (Gibco) containing 0.1mM non-essential amino acids (NEAA) (Gibco), 2mM Glutamine (Gibco) and 1mM Sodium pyruvate (Gibco), 100u/ml Penicillin, 0.1mM 2-mercaptoethanol (Sigma) and 10% Fetal Calf Serum (F2442, Sigma) and 1000U/ml ESGRO (Sigma) until their differentiation. ESC were transitioned towards EpiLCs according to previously published protocols (Hayashi et al., 2011). Briefly 150,000 EpiLC were plated in a 12-well plate coated with human plasma fibronectin (16.7ng/ml) in N2B27 medium containing Activin A (20ng/ml, Peprotech), bFGF (12ng/ml, R&D Systems) and 1% KSR (Gibco). Cells for IF were plated in 8-well chamber slide permanox plastic (ThemoFisher) coated with human plasma fibronectin. The medium was changed every day. EpiLC were analyzed at the time points indicated in the figures. Pharmacological inhibitors were added to the concentrations indicated. Dorsomorphin and SB431542 were obtained from Tocris.

### Immunostaining on EpiLCs

Cells were fixed for 10 minutes in 4% PFA at RT followed by 4 washes with PBS, 10 minutes each. Immunostaining was performed in blocking buffer. The following primary antibodies were incubated overnight in blocking buffer: CDH1 (1:500 Sigma-Aldrich), CDH2 (1:300, Cell Signaling Technology), T (1:400, Cell Signaling) or T(1:500 R&D systems), KDR (1:50, BD Pharmigen), pSMAD1/5/9 (1:100 Cell Signaling), FOXA2 1:500 (Abcam), SOX2 (1:400 Cell Signaling), POU5F1 (1:500, Santa Cruz Biotech.), NANOG (1:300 BD Biosciences). Fluorescent-secondary antibodies (Invitrogen) were diluted at 1:400 in blocking buffer and incubated for 1 hour at RT (room temperature). Cells were mounted in ProLong Gold (ThermoFisher) and imaged using a Leica-upright SP5 laser point-scanning microscope or a Zeiss LSM 880 laser-scanning confocal microscope. Confocal images were reconstructed and analyzed by using Volocity software package (PerkinElmer).

### RT-qPCR analysis

EpiLC were lysed using in RLT Lysis buffer (RNeasy, Qiagen) at the time points indicated. RNA from cells was extracted using the RNeasy Mini Qiagen kit (Qiagen). 500ng of RNA from EpiLC was reverse transcribed to cDNA using Superscript III-RT (Thermo) and random hexamers and the Qiagen kit. cDNA from EpiLC was diluted at concentrations of 1:10, 1:5 and 1:3. qPCR was performed using Power Up SYBR Green Master Mix (Applied Biosystems). Each sample was analyzed in triplicate, samples were normalized using GAPDH and the differential expression was obtained using the delta-delta CT method. The following oligonucleotide primers were used:

GAPDH Forward: GGGTCCCAGCTTAGGTTCAT, GAPDH Reverse: CGTTGATGGCAACAATCTCT; T Forward: CTCGGATTCACATCGTGAGAG, T Reverse: AAGGCTTTAGCAAATGGGTTGTA; KDR Forward: CGAGACCATTGAAGTGACTTGCC, KDR Reverse: TTCCTCACCCTGCGGATAGTCA; FOXA2 Forward: CGAGCACCATTACGCCTTCAAC, FOXA2 Reverse: AGTGCATGACCTGTTCGTAGGC

## ACKNOWLEDGEMENTS

We thank Frank Costantini (Axin2 mutants) for mice, members of the Anderson and Hadjantonakis labs for discussions and feedback on the work, the Memorial Sloan Kettering Cancer Center (MSKCC) Mouse Genetics Core Facility, Integrated Genomics Operation for bulk and single-cell sequencing, and the Bioinformatics Core facility for analysis of bulk RNAseq data. MSKCC’s core facilities are supported by an NCI Cancer Center Core Grant (P30CA008748). This study was supported by the NIH, R01HD094868 (KVA and AKH), R01DK127821 (AKH), and a Collaborative Award from The Wellcome Trust (BG and AKH). LTGH was funded by a Wellcome Early Career Award (226309/Z/22/Z). RHM was supported by a postdoctoral fellowship from Pew Latin American Fellows Program for part of this work.

## AUTHOR CONTRIBUTIONS

RMH and KVA conceived the project; RHM performed immunofluorescence, mRNA *in situ* and bulk RNAseq analyses of mouse embryos, and pluripotent stem cells; RHM, SN and YYK generated scRNAseq libraries from embryos; SN, LTGH, BT and AKH processed and analyzed scRNAseq data; BG, EL, AKH and KVA procured funding and provided project supervision; RMH wrote a draft of the manuscript with input from EL and AKH; all authors provided input and edited the manuscript prior to submission.

## SUPPLEMENTARY FIGURE LEGENDS

**Figure S1.** (**A**) Single optical sections of whole-mount immunofluorescence detecting T and CDH1 in E5.75 wild-type, *Axin1^fus/fus^;Axin2^-/-^* and *Smad2^-/-^*embryos. (**B**) Brightfield images of E9.5 wild-type and *Axin1^ΔEpi^;Axin2^+/+^* mutant embryos. Scale bar is 100μm. (**C**) IF on E6.5 longitudinal sections for SOX2, NANOG and POU5F1 in wild-type, *Axin1^ΔEpi^;Axin2* and *Smad2* mutant embryos. (**D**) Graph showing ratio of T and CDH2-positive cells respect to total nuclei on E6.5 wild-type and *Smad2^-/-^* mutant embryos. Bars represent the mean± SEM. Statistically significant differences were determined by t-test. ***p<0.0008.

**Figure S2.** (**A**) UMAP from a down sampled version of the gastrulation atlas that spans E6.5 to E8.5. Cells are colored by the extended gastrulation atlas cell type label in **A** and embryonic stage in **B. (C)** Dot plot showing log normalized expression levels of top marker genes (avg_log2FC > 0.5) for the cell types show in Figure 1A from the control and *Axin1^ΔEpi^;Axin2^-/-^* datasets. The size of the dot indicates the percentage of cells expressing the marker gene. (**D**) Heat map showing top differentially genes upregulated (red) and downregulated (green) in the E6.5 single embryo (se) bulk transcriptome of wild-type and *Axin1^ΔEpi^;Axin2^-/-^* mutant embryos. (D) Graph showing upregulated genes in the E6.5 se-transcriptomes of *Axin1^ΔEpi^;Axin2* mutant embryos associated with differentiation of blood precursors. (**F**) Graph showing ExM genes upregulated in E6.5 se-transcriptomes of *Axin1^ΔEpi^;Axin2* mutant embryos.

**Figure S3.** (**A**) Graph showing the ratio of KDR, CDX2 and FOXF1-positive cells relative to the total nuclei on E6.5 wild-type, *Axin1^ΔEpi^;Axin2* and *Smad2* mutant embryos. Bars represent the mean± SEM. Statistically significant differences were determined by t-test. *p<0.01, **p<0.004, ***p<0.0001. (**B**) Graph showing se-bulk RNA expression levels for BMP ligands in the transcriptome of E6.5 *Axin1^ΔEpi^;Axin2* double mutant embryos.

**Figure S4.** (**A**) Heat map showing top differentially genes upregulated (red) and downregulated (green) in the se-transcriptome of E7.5 wild-type and *Axin1^ΔEpi^;Axin2^-/-^*mutant embryos. (**B**) Gene Ontology terms downregulated (blue) and upregulated (salmon) in the E7.5 se-transcriptome of *Axin1^ΔEpi^;Axin2* mutant embryos respect to wild-type. (**C**) IF for KDR, CDX2 and FOXF1 on longitudinal sections of E7.5 wild-type, *Axin1^ΔEpi^;Axin2^-/-^* and *Smad2^-/-^* embryos. (**D**) Graph showing bulk RNA expression levels of NODAL target genes downregulated in *Axin1^ΔEpi^;Axin2* mutant embryos with respect to wild-type at E6.5 and (**E**) E7.5.

**Figure S5.** (**A**) IF detecting SOX2, NANOG and POU5F1 in wild-type and *Axin1^fus/fus^;Axin2^-/-^*ESCs grown in 2i conditions. (**B**) IF detecting T on wild-type and *Axin1;Axin2* EpiLC mutant cells after differentiating for 24 and 48 hours. (**C**) IFs for EMT markers CDH1, T and CDH2 in wild-type, *Axin1*/*Axin2*and *Smad2* mutant EpiLC after differentiating for 48 hours. Cartoon on top is describing the experimental strategy. “Created with BioRender.com”.

**Table S1.** Genes upregulated in the se-bulk RNA transcriptome of E6.5 control and *Axin1^ΔEpi^;Axin2^-/-^* mutant embryos p<0.05.

**Table S2.** Genes downregulated in the se-bulk RNA transcriptome of E6.5 control and *Axin1^ΔEpi^;Axin2^-/-^* mutant embryos p<0.05.

**Table S3.** Genes upregulated in the se-bulk RNA transcriptome of E7.5 control and *Axin1^ΔEpi^;Axin2^-/-^* mutant embryos p<0.05.

**Table S4.** Genes downregulated in the se-bulk RNA transcriptome of E7.5 control and *Axin1^ΔEpi^;Axin2^-/-^* mutant embryos p<0.05.

