## Supplementary figures and images for "Axin1 and Axin2 regulate the WNT-signaling landscape to promote distinct mesoderm programs"

### Figure S1, Figure S2, Figure S3, Figure S4, Figure S5

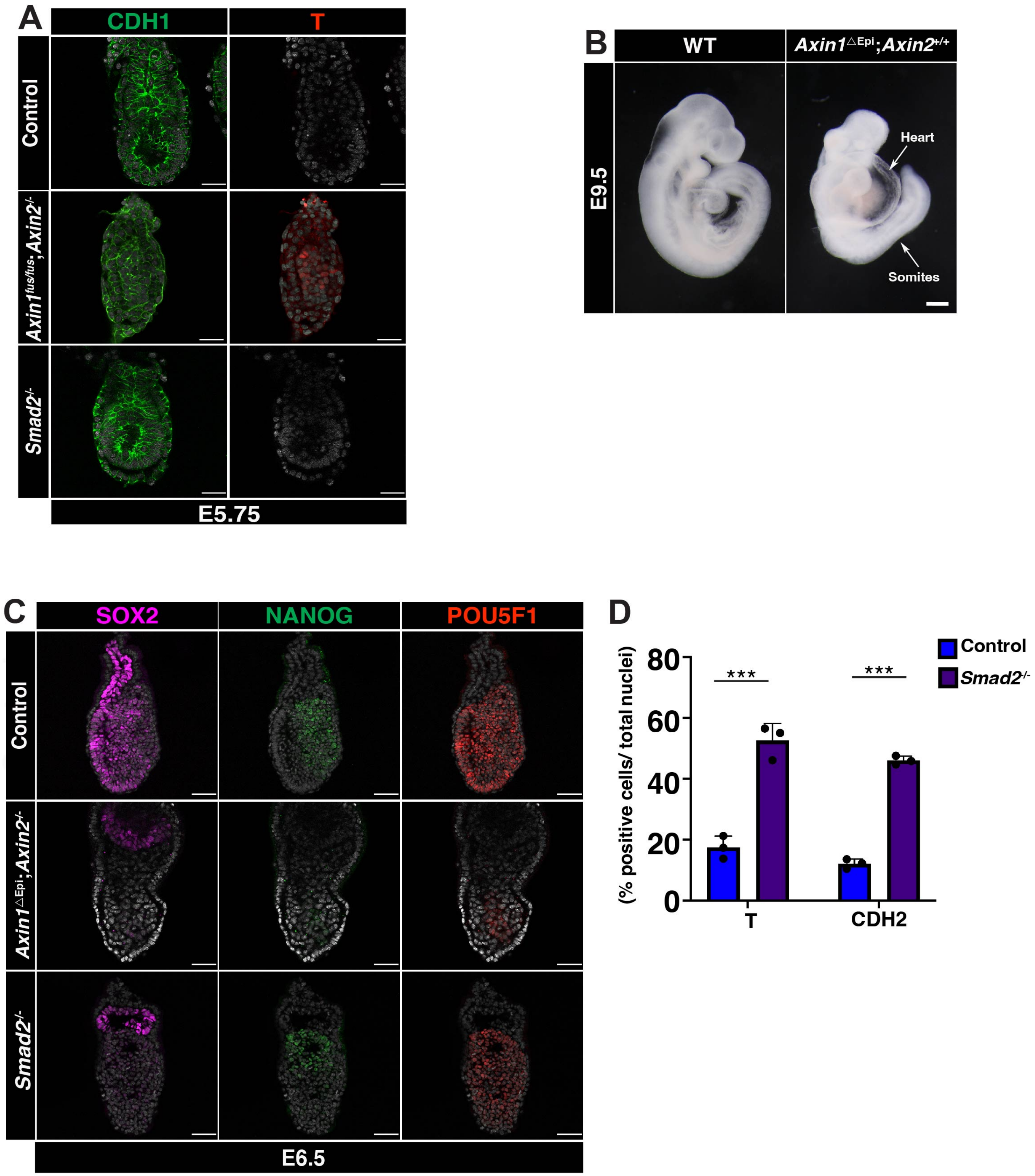

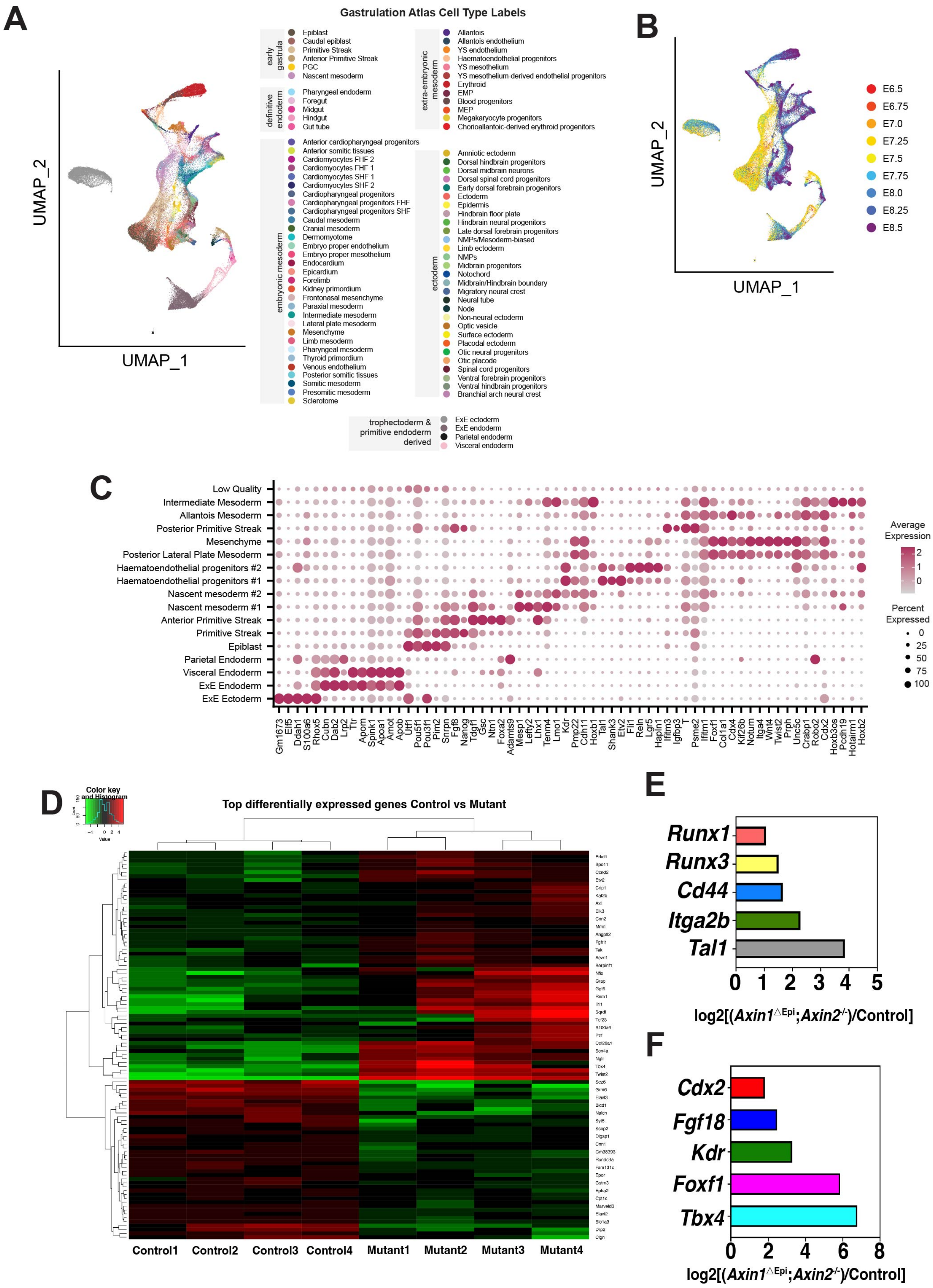

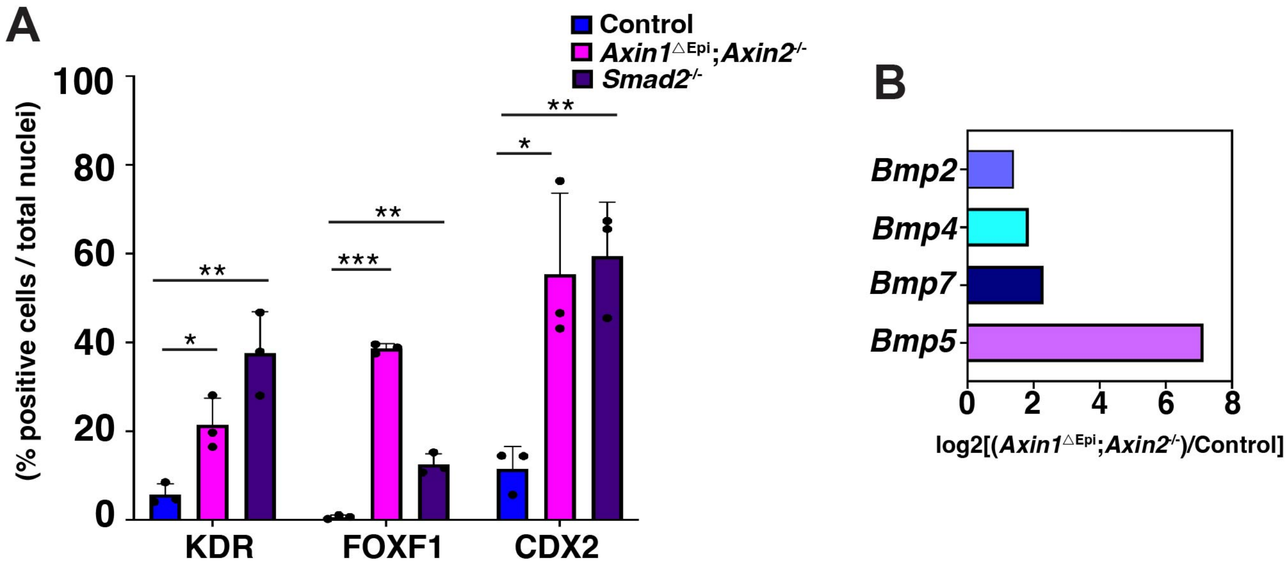

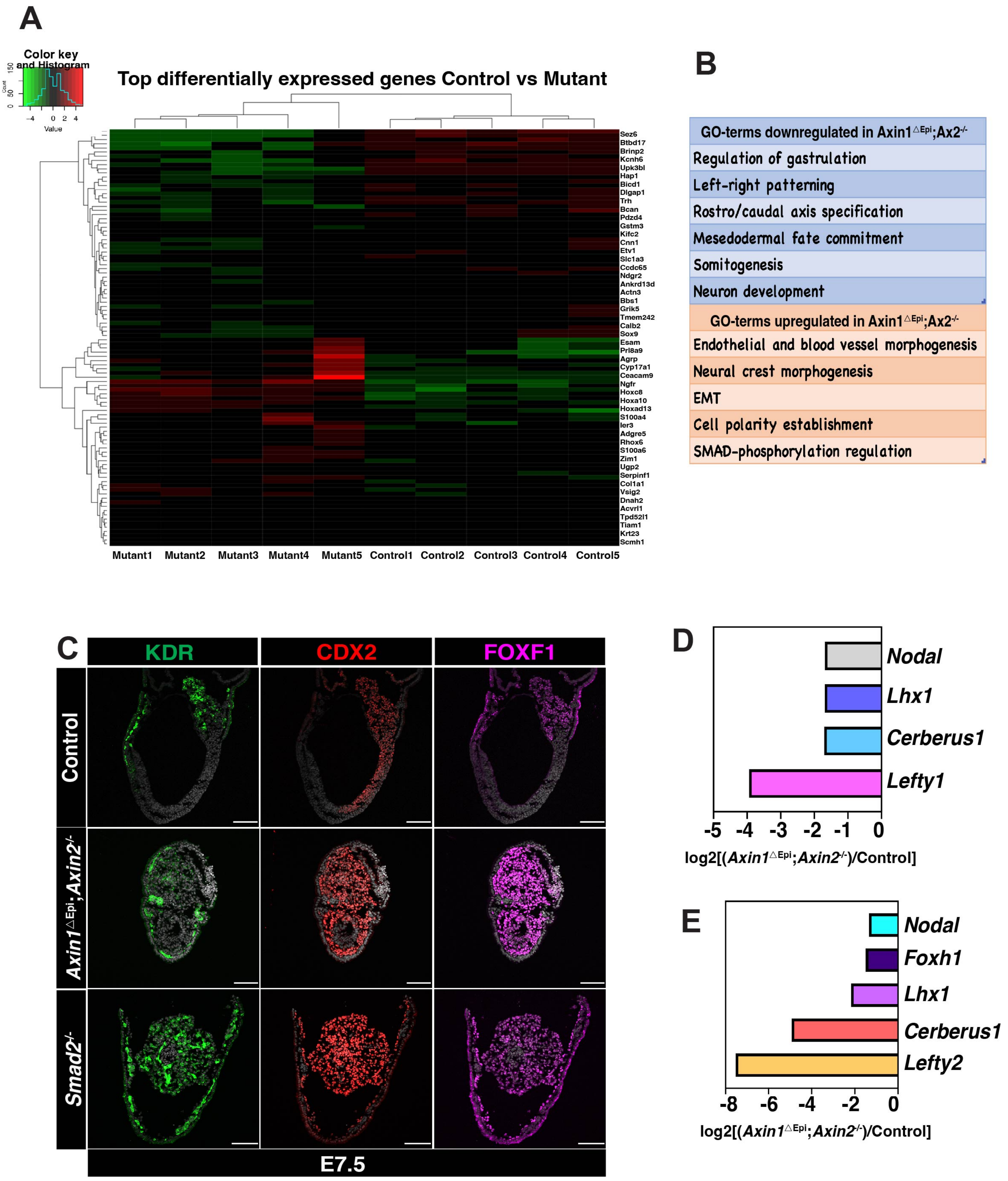

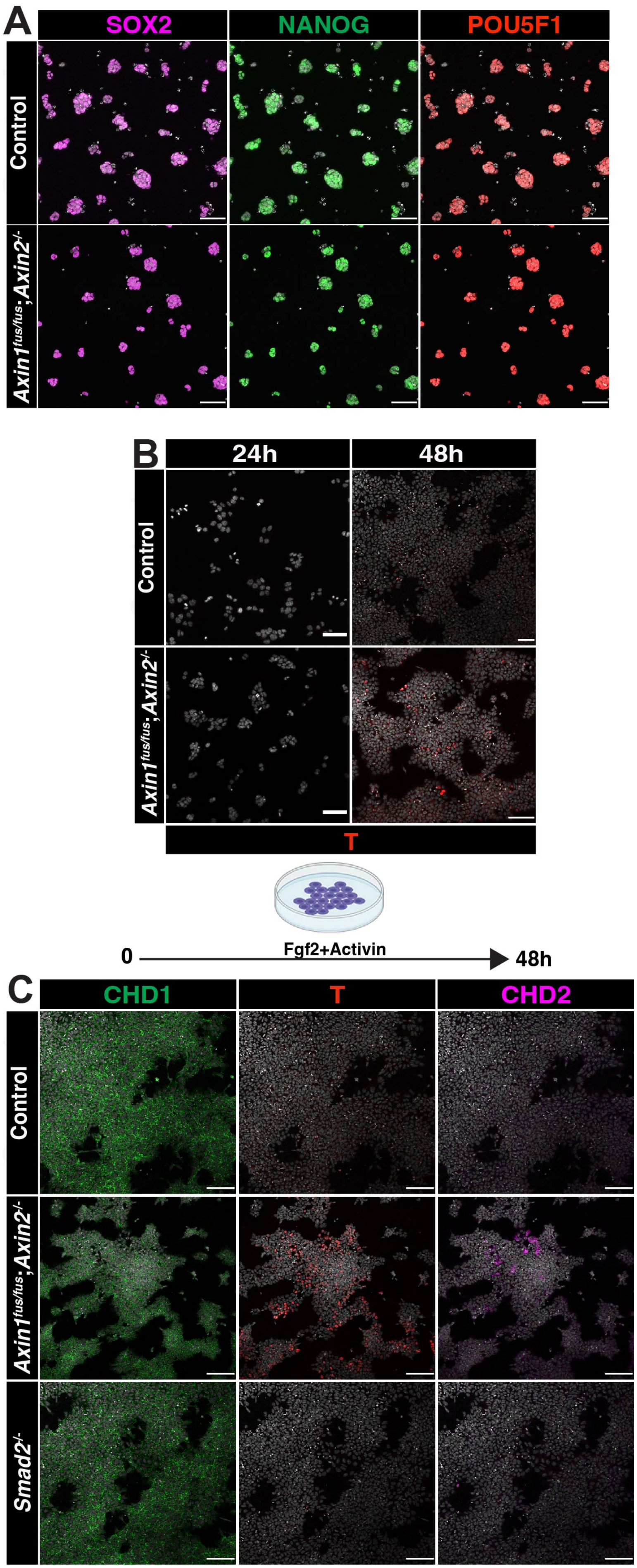
